# The human CD8 T stem cell-like memory phenotype appears in the acute phase in Yellow Fever virus vaccination

**DOI:** 10.1101/808774

**Authors:** Silvia A. Fuertes Marraco, Amandine Bovay, Sina Nassiri, Hélène Maby-El Hajjami, Hajer Ouertatani-Sakouhi, Werner Held, Daniel E. Speiser

**Author notes:** Correspondence : Silvia. A Fuertes Marraco, Département d’Oncologie CHUV-UNIL, AA-82, Biopôle 3, Centre de Laboratoires d’Epalinges, Chemin des Boveresses 155, CH-1066 Epalinges.

## Abstract

Long-term memory is a fundamental feature of cytotoxic CD8 T cell immunity. Yet when do memory cells arise, especially in humans, is poorly documented, the pathways of effector / memory cell differentiation being largely debated. Based on a cross-sectional study, we previously reported that the live-attenuated Yellow Fever virus vaccine YF-17D induces a stem cell-like memory (SCM) CD8 T cell population persisting for at least 25 years. Here we present longitudinal data revealing that an activated SCM phenotype is distinctly detectable early on following YF-17D vaccination, i.e. at the same time as activated effector cells. In the long-run, the cells that express the transcription factor T cell factor 1 (TCF1) preferentially persist, consistent with the role of TCF1 in memory establishment. By performing t-distributed Stochastic Neighbour Embedding of flow cytometry data on standard differentiation and activation markers, we obtained a time-lapse representation of the dynamics of the CD8 T cell response: SCM cells appear early and remain closely related to the baseline Naive cells, while effector cells burst out of baseline and gradually contract after the peak of the response. Altogether, we observe heterogeneity in differentiation phenotypes in both the acute phase and the decade-long-term phases, including cells with memory phenotypes very early in the response. As opposed to models where memory cells develop from effector cells, our data support differentiation models where long-term memory cells are established by the early decision to retain proximity to the Naive state in a memory-dedicated pool of cells.

## Introduction

The capacity to remember a pathogen and effectively protect the organism against it long-term is a fundamental property of the adaptive immune response. This is also relevant for tumour immunology since it is now well established that strong and long-lasting cytotoxic CD8 T cell responses correlate with better prognosis for cancer patients (Fridman et al., 2012), and that innovative immunotherapies can defeat various types of metastatic cancers with unprecedented long-term success (Ribas and Wolchok, 2018).

Once the naïve T cells are primed upon antigenic encounter, the various functionalities of CD8 T cells are ensured by a heterogeneity of cells, with varying degrees of memory and effector functions. Initially classified into only two functional types (effector or memory), the heterogeneity of CD8 T cells has been more comprehensively defined over the last decade. The venue of transcriptomic and epigenetic profiling complementing functional assays has revealed a continuum of phenotypes with varying longevity, self-renewal, proliferative potential, expression of homing, costimulatory and transcription factors, and functions including cytokine secretion and cytotoxicity (Crompton et al., 2015; Farber et al., 2013; Gattinoni et al., 2012; Gray et al., 2014; Roychoudhuri et al., 2015). Globally, effector cells display cytotoxicity and readily produce cytokines, while memory cells resemble more the Naïve cells based on their high proliferative capacity and potential to generate effector progeny, together with long-term persistence and self-renewal (so called stemness). Traditionally, surface markers (including distinct homing molecules) and transcription factors have been used to define the various CD8 T cell subsets. In humans, classic subsets are primarily identified on the basis of surface C-C motif chemokine receptor 7 (CCR7) and CD45RA expression, with Naïve being CCR7+ CD45RA+, the central memory (CM) being CCR7+ CD45RA-, and the CCR7- effector memory subsets split into CD45RA- effector memory (EM) and effector memory CD45RA+ (EMRA) cells (Sallusto et al., 2004). More recently, the stem cell-like memory (SCM) subset was revealed among CCR7+ CD45RA+ cells (within the classic Naïve gate) on the basis of positive expression of markers such as CD58, CXCR3, IL2Rb and the more prominently used CD95 marker (Gattinoni et al., 2011; Lugli et al., 2012).

Along with the increasingly comprehensive characterization of the heterogeneity of CD8 T cell phenotypes and functions, several models have emerged to describe the differentiation pathways of antigen-experienced CD8 T cells, addressing the genealogy of memory and effector cells. The initially proposed model of CD8 T cell differentiation is linear: it suggests a sequential differentiation of naïve cells, first into effectors that predominate in the acute phase, followed by differentiation of a fraction of effector cells into memory cells, as the response contracts and the remaining effector cells die out or become terminally senescent. In the mouse system, the linear model evolved to describe early effector cells (EEC) that give rise to two types of effector cells: one Short-lived effector cell (SLEC) and another Memory precursor effector cell (MPEC) – long-term memory thus predominantly originates from MPECs (Crauste et al., 2017; Lefrançois and Obar, 2010; Yuzefpolskiy et al., 2014). Alternative models have proposed that memory cells diverge from effector cell differentiation, without an obligatory acute effector stage. For instance, the so-called bifurcative model proposes an immediate divergence from the naïve cell: in a first asymmetric cell division, the antigen-primed naïve cell splits into distinct daughter cells, each with a distinct memory or effector fate (Ahmed et al., 2009; Moulton and Farber, 2006). More recently, the proposed models integrate the large and gradual heterogeneity of memory and effector cells, based on the observed continuum of whole transcriptome and epigenetic profiles (Crompton et al., 2015; Henning et al., 2018; Restifo and Gattinoni, 2013). These suggest that CD8 T cells may undergo progressive differentiation, from the naïve, to the SCM, CM, EM and EMRA cell stages, and all the various subsets may give rise to effector progeny or show effector function in their activated state (Farber et al., 2013; Mahnke et al., 2013).

To date, it is still controversial whether memory results from an early decision to diverge from effector fate or whether a fraction of effectors gives rise to memory cells. Recent studies continue debating whether long-lived memory cells display an epigenetic imprint that would correspond to an effector phenotype past (Akondy et al., 2017; Ben Youngblood et al., 2017) or whether stemness (highest memory potential) is preserved epigenetically in antigen-primed naïve cells that become memory cells and it is the silencing of memory / stemness genes that drives effector differentiation (Pace et al., 2018). One limitation is that nascent memory cells are not easily detectable or may not be distinguished from effector cells in the activated, acute phase (Opata and Stephens, 2013). Markers such as IL7Ra have been highlighted to identify the precursors of long-lived memory cells (the MPECs in mice), distinct from the majority of activated cells that die after the acute phase (Kaech and Cui, 2012). One major factor that is essential to sustain memory formation is the transcription factor T cell factor 1 (TCF1, encoded by the *TCF7*/*Tcf7* gene) (Jeannet et al., 2010; Utzschneider et al., 2016; Zhao et al., 2010; Zhou et al., 2010). TCF1 is expressed at high levels in Naïve and memory but not in effector cells (Kratchmarov et al., 2018; Roychoudhuri et al., 2015; Willinger et al., 2006), and is epigenetically regulated during CD8 T cell differentiation (Abdelsamed et al., 2017; Crompton et al., 2015) as one major gene involved in effector differentiation arrest and maintenance of stemness (Gattinoni et al., 2009; Pace et al., 2018; Wu et al., 2016). Recently, we showed that inflammatory cytokines suppress TCF1 and facilitate effector differentiation (Danilo et al., 2018).

Overall, studies on the identification of precursors and discernment of early fate decisions rely on genetic manipulation and the adoptive transfer or deletion of cells to test their progeny potential, which is limited to mouse models. Yet a major level of complexity in the study of CD8 T cell differentiation is the idiosyncrasies in mouse *versus* human systems. While fundamental phenomena may be shared, in practice, there are basic differences in the markers used to classify CD8 T cell subsets. Therefore, and in complement to the ontological questions that can readily be addressed in the mouse experimental system, the evidence that originates from the study of human CD8 T cells is uniquely valuable. One human model that has been particularly informative to fully apprehend optimal immunogenicity in humans, including the study of CD8 T cell differentiation, is the acute response to the live-attenuated Yellow Fever virus vaccine YF-17D (Miller et al., 2008; Pulendran et al., 2013). We previously found that YF-17D vaccination induces a population of SCM cells and showed that these memory cells last for decades representing the most stably persisting T cell population ever described (Fuertes Marraco et al., 2015; Marraco et al., 2015). However, the earliest time-point after vaccination that we studied was 3.6 months, well after the acute phase of the response. There is currently no information on when SCM cells appear during an immune response in humans. Here, we aimed to study the distribution and dynamics of human CD8 T cell subsets during the first few days to months after YF-17D vaccination based on a longitudinal clinical study protocol (i.e. including the acute phase of the response), combined with analysis in the decade-long-term based on our previous cross-sectional study cohort.

## Materials and Methods

### Study design, population and ethics statement

Samples used in this study originated from peripheral blood of healthy volunteers aged 18 to 65 years that participated in one of two study protocols on YF-17D vaccination (Stamaril, Sanofi Pasteur). Donors from the first cohort “YF1” (study protocol 329/12) had a history of YF-17D vaccination ranging from 3.6 months to 23.74 years (cross-sectional) and donated blood in the local Blood Transfusion Center (Service régional vaudois de transfusion sanguine, 1066 Epalinges), as we described previously ((Fuertes Marraco et al., 2015). Donors from the second cohort “YF2” (study protocol 324/13) were in the prospect of receiving the YF-17D vaccine in view of travelling to endemic areas and participated to longitudinal sampling before and several time-points after YF-17D vaccination, in collaboration with the local vaccine center Centre de vaccination et de médecine des voyages (Policlinique Médicale Universitaire (PMU), Lausanne). The full metadata details of the two cohorts are listed in Table S1. The study protocols were approved by the Swiss Ethics Committee on research involving humans of the Canton of Vaud (CH). All participants provided written informed consent.

### Peripheral blood collection and processing

Peripheral blood samples were collected and immediately processed for cryopreservation awaiting experimental use. Peripheral blood mononuclear cells (PBMC) were obtained from anti-coagulated whole blood diluted 1:1 in phosphate buffered saline (PBS) and overlaid on Lymphoprep for density gradient fractionation (30 min at 400g without break) and were cryopreserved in complete RPMI 1640 supplemented with 40% fetal calf serum (FCS) and 10% dimethyl sulfoxide. Plasma samples were obtained from the supernatant of EDTA-coated blood tubes after centrifugation at 1’000g for 15 min at RT followed by a second centrifugation at 8’000g for 10 min at 4°C.

### Assay to determine copy numbers of the Yellow Fever virus YF-17D

Yellow Fever virus (YFV) load was quantified using 1ml of plasma from EDTA-anticoagulated blood based on a Taqman Real-time PCR assay to detect YFV genome copies as previously described (Akondy et al., 2015).

### Flow cytometry staining, acquisition and analysis

On the day of the experiment, frozen vials of PBMC were thawed in RMPI containing 10 μg / ml of DNAse I (Sigma) and resuspended in fluorescence-activated cell sorting buffer (FACS buffer: PBS with 5mM EDTA, 0.2% Bovine Serum Albumin and 0.2% sodium azide). Thawed PBMC were subjected to CD8+ T cell selection using the negative enrichment kit from Stem Cell. CD8 T cell-enriched samples were then stained for flow cytometry according to target panels and cytometers as summarized in Table S2 and with reagents as listed in Table S3. Stainings were made in sequence depending on the target, as follows: 1) first, cells were stained with multimers for 30 min at 4°C in FACS buffer and washed in FACS buffer, 2) surface antibodies were added in FACS buffer and washed with PBS prior to 3) staining with fixable viability dye in PBS and washed with PBS, 4) cells were then fixed overnight at 4°C and washed in permeabilization buffer before 5) intracellular staining in permeabilization buffer at 4°C (the primary rabbit anti-TCF1 and the secondary fluorochrome-conjugated anti-rabbit IgG were stained in two subsequent steps). The fixation and permeabilization buffers were from the Foxp3 staining kit from eBioscience. Washes were made by centrifugation at 450g for 7 min. Samples were resuspended in FACS buffer for acquisition. For samples in the YF2 study, the baseline sample vial originally contained 1.5 x 10e7 frozen PBMC and the remainder of time-points’ vials contained 10e7 frozen PBMC – the complete volume of stained samples was finally acquired. Cytometers were the Gallios (Beckman Coulter, 3 laser, 10-color) and the LSR II Special Order Research Product (Beckton Dickinson, 5 laser including UV, 13- or 14- color). Before each acquisition, the cytometer setup and tracking (CST) was ran in order to normalize channel voltages across experiments using the same instrument configuration and experimental layout. Flow cytometry FCS data files were analyzed in FlowJo 9.7.7, except for the analyses using t-distributed Stochastic Neighbor Embedding (tSNE) for which the corresponding plugin in FlowJo 10.4.2 was used. Downsampling, concatenation or exports of specific populations and samples were performed as indicated in the figure legends also in FlowJo 10.4.2. For the longitudinal tSNE analyses of A2/LLW-specific CD8 T cells, all single live A2/LLW-specific CD8 T cell events of the longitudinal series were concatenated and thus are represented in proportions corresponding to the original numbers of PBMC thawed, which are equal across time-points (corresponding to 10e7 PBMC thawed) except for the baseline which is 1.5-fold larger (1.5 x 10e7 PBMC thawed). The detection threshold for multimer positive populations was 0.01% of total CD8 T cells and at least 10 events (horizontal dotted line in Figure 1C and D), based on control stainings using HLA-A*02 negative samples stained with tetramer and unstained controls. The positivity threshold for each marker was set according to distinct negative and positive populations in bulk CD8 T in resting and/or activated samples; for the indirect TCF1 staining, the negative signal was further validated with secondary antibody-only controls.

**Figure 1.**
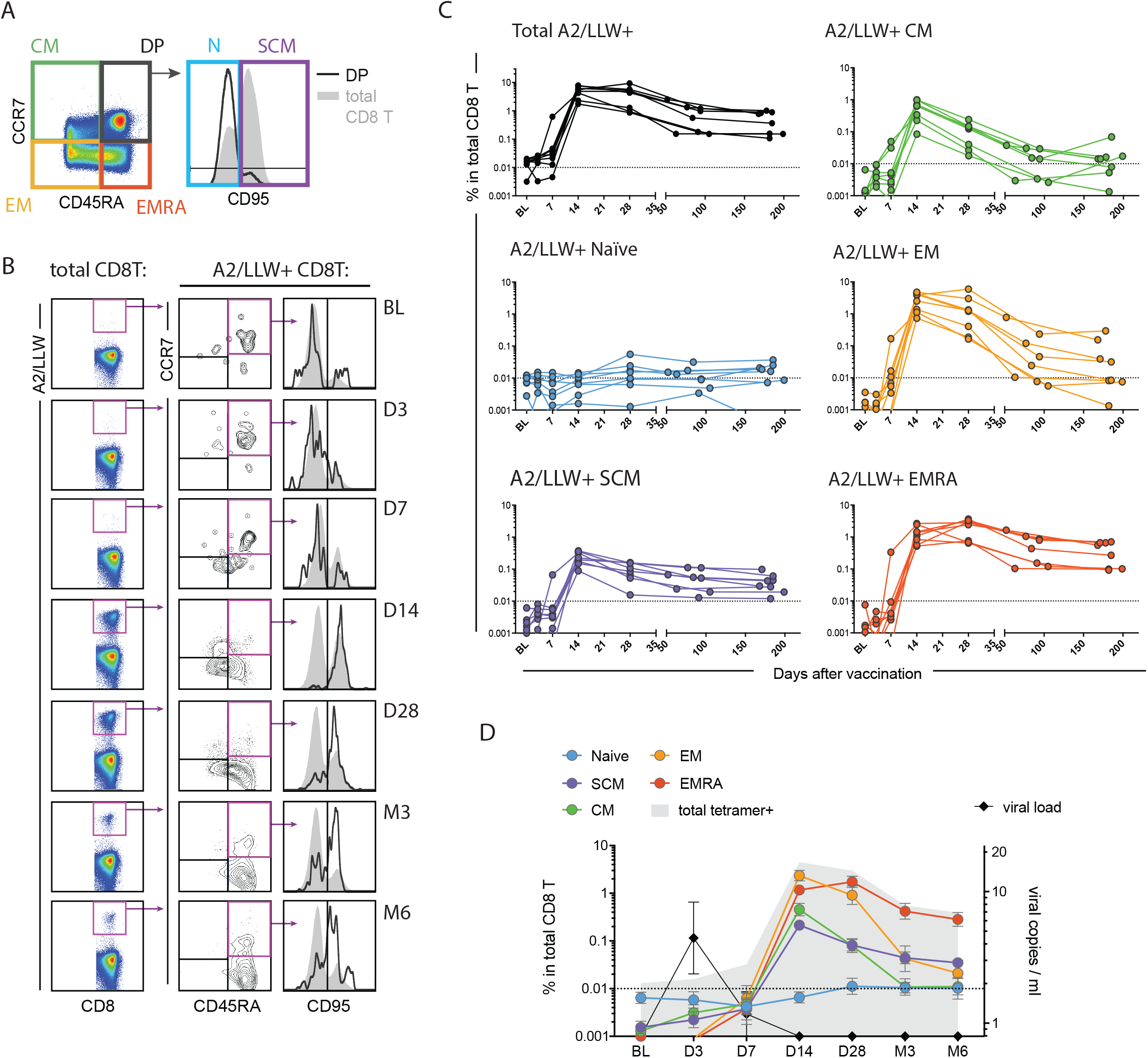
Quantification of A2/LLW-specific CD8 T cell subsets during the first six months after YF-17D vaccination. **(A)** Flow cytometry gating strategy to define CD8 T cell subsets: Central memory (CM: CCR7+ CD45RA-), Effector memory (EM: CCR7- CD45RA-) and Effector memory CD45RA+ (EMRA: CCR7- CD45RA+) cells; CCR7 and CD45RA double positive (DP) cells are further subdivided into Naïve (CD95-) and stem cell-like (SCM, CD95+) subsets. **(B)** Flow cytometry analysis of A2/LLW tetramer+ CD8 T cells, showing the 7 longitudinal time-points of the representative donor (LAU 5089): CD8+ A2/LLW tetramer+ cells were analyzed for subset distribution as in A. **(C) and (D)** Quantification of the frequency of A2/LLW tetramer+ within total peripheral CD8 T cells, in N=8 donors with 7 longitudinal time-points. In C, frequencies are shown for total A2/LLW+ or per A2/LLW+ subset, and each donor is line-connected across its dotted time-points. In D, the data is pooled showing average and standard error of the mean (N=8) per population as indicated (left y-axis); viral load data is complemented (right y-axis). The dotted line in C and D indicates the multimer detection threshold of 0.01% of total CD8 T cells. Time-points are BL: baseline, D3: Day 3, D7: Day 7, D14: Day 14, D28: Day 28, M3: circa 3 months, M6: circa 6 months (Table S1 shows full details of the cohort).

### Quantifications and statistical analyses

Flow cytometry data analyzed with FlowJo was quantified based on tabulated exports of the frequencies and events in the gates of interest. Calculations and data display thereafter was performed using the softwares Microsoft Excel 15.21.1, GraphPad prism 7.0c and SPICE v5.35 (for co-expression analyses). Statistical values were obtained as detailed in each figure legend (on the basis of normality tests), where trend = p>0.05 and <0.10, * = p<0.05, ** = p<0.01, *** = p<0.001 and ns = not significant. For the SPICE analyses, p-values originate from the built-in t-test in SPICE using 10’000 permutations. Longitudinal modeling of the flow cytometry data was achieved using linear mixed effects splines. In brief, linear splines with 3 internal knots and a random intercept was fit using the lme4 package in R (Bates et al., 2015). Pairwise comparison of fits to individual subsets was performed by fitting a null model to pooled data from the two subsets, a full model with distinct fits capturing the trends in each subset, and using the likelihood ratio test to assess the difference between these two nested models using the Chi square distribution. Resulting p-values were further adjusted for multiple comparisons using the Bonferroni method.

## Results

### CD8 T cells with a CCR7+ memory phenotype expand in the acute phase of YF-17D vaccination

In order to study the early dynamics of CD8 T cell differentiation, we recruited healthy volunteers that were going to receive the YF-17D vaccine in order to obtain peripheral blood samples before and at several time-points after vaccination (including early days and up to 6 months after vaccination). The study schedule and cohort are detailed in Table S1A: “Longitudinal cohort: YF2 study”. Using peptide-MHC multimers, we detected CD8 T cells specific for the immunodominant HLA-A*02-restricted epitope of the Non-Structural 4b protein of Yellow Fever virus (the LLWNGPMAV epitope (Akondy et al., 2009; Blom et al., 2013; de Melo et al., 2013)), hereafter referred to as “A2/LLW”) in eight HLA-A*02+ vaccinees. The phenotypes of CD8 T cell differentiation were determined based on the classic markers CCR7 and CD45RA (Sallusto et al., 2004) as shown in Figure 1A to detect Central Memory (CM: CCR7+ CD45RA-), Effector Memory (EM: CCR7- CD45RA-) and Effector Memory CD45RA+ (EMRA: CCR7- CD45RA+); within the CCR7 CD45RA double-positive gate, Naïve and Stem Cell-like Memory (SCM) subsets were discriminated based on CD95 expression (Gattinoni et al., 2011; Lugli et al., 2012; Mahnke et al., 2013). Of note, the aforementioned subset nomenclature describes resting human CD8 T cells; for the purpose of longitudinal consistency we maintain this nomenclature yet we highlight that acutely activated human effector cells downregulate CCR7 and phenotypically coincide with EM and EMRA (Mahnke et al., 2013).

As previously described (Akondy et al., 2009), we observed massive expansion of A2/LLW-specific CD8 T cells with a peak around day 14 post-vaccination, with largely predominant CCR7- phenotypes (Figure 1B and C: “Total A2/LLW+”, “EM” and “EMRA” plots). Remarkably, detailed longitudinal quantification also showed expansion of CCR7+ memory phenotype cells: both CM and SCM cells were clearly detected and expanded by day 14 (Figure 1C and D). After the peak at day 14, EM cells contracted, while EMRA cells continued to increase slightly until day 28. At the later time-points, especially by 6 months, it was evident that EMRA and SCM subsets persisted, while EM and CM subsets continued to fade away. This later observation is in line with our previous report where the EMRA and SCM subsets were the two subsets predominantly detected in the long-term (range of years to decades), the SCM cells being the most stable memory cell subset described so far (Fuertes Marraco et al., 2015).

In addition, we determined whether CCR7+ memory phenotype cells detected during the early phase post-vaccination co-existed with antigen, i.e. before viral clearance. Live-attenuated vaccine virus YF-17D was detectable at days 3 and/or 7 in the plasma of five out of the eight vaccinees (Figure 1D and Figure S1). The analysis was challenged by the fact that A2/LLW multimer positive cells were close to the limit of detection at baseline and at days 3 and 7 (only samples that showed total A2/LLW+ CD8 T cells superior to 0.01% of total CD8 T were further analyzed for subset distribution). In two donors, CM (in donor LAU 5088) and both CM and SCM (in donor LAU 5080) cells were detected at the same time-point when virus was detectable (Figure S1). These data show that cells with a memory phenotype can arise before antigen is cleared, well ahead of the contraction phase of the response.

### SCM and CM phenotype cells are activated at the peak of the response

In parallel to the rise in frequencies, the acute phase of the T cell response is characterized by the expression of activation markers as previously described in total A2/LLW+ CD8 T cells (Akondy et al., 2009; Blom et al., 2013; Querec et al., 2009). In order to address how activation status compared across CD8 T cell subsets, we measured the longitudinal expression of activation markers: CD69, CD38, HLA-DR and PD1, within each subset. At the peak of the response (day 14), the analyses clearly showed that the CCR7+ memory subsets (SCM and CM) were extensively activated, in fact as much as the CCR7- EM and EMRA subsets (Figure 2). The early activation marker CD69 was most highly expressed at days 3 and 7, while HLA-DR, CD38 and PD1 peaked at day 14 (Figure S2). Beyond day 14, CD38 clearly diminished while HLA-DR and PD1 partially persisted (Figure S2).

**Figure 2.**
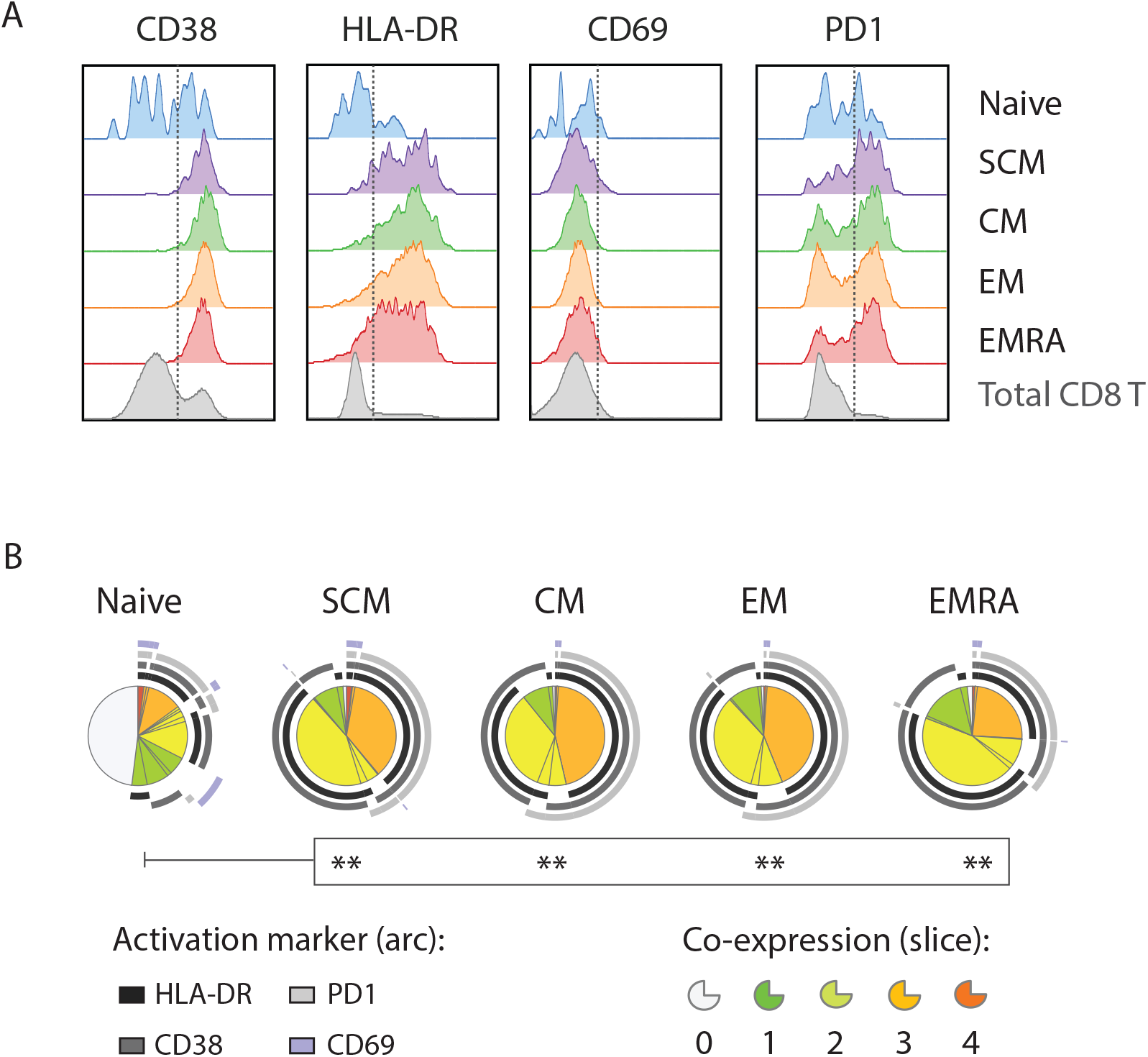
Activation of A2/LLW-specific CD8 T cell subsets at the peak of the response. **(A)** Flow cytometry profiles at day 14 showing each activation marker and subset, as indicated; total CD8 T cells are shown as a reference. The data are from donor LAU 5089. **(B** Pie charts showing frequencies of the combinatorial expression of the four indicated activation markers, per subset: each arc designates one marker, each slice a number of markers co-expressed (representation based on SPICE software). N=7 donors were analyzed at day 14 post-vaccination and only detectable populations were quantified: in Naïve (n=4/7), SCM (n=7/7), CM (n=7/7), EM (n=7/7), EMRA (n=7/7). P-values (built-in t-test in SPICE): ns = not significant, * < 0.05, ** < 0.01, *** < 0.001.

Of note, in the aforementioned longitudinal analyses, we observed that A2/LLW+ CD8 T cells were still present in the Naïve gate (as defined by CCR7+ CD45RA+ CD95-) after vaccination and that they remained relatively stable over time (Figure 1). Interestingly, these post-vaccination Naïve cells did show substantial activation at the peak of the response (Figure 2 and S2). The nature of these Naïve-gated cells will be discussed.

### TCF1+ CD8 T cells preferentially persist for decades

Given the central function of TCF1 in memory establishment (Gattinoni et al., 2009; Jeannet et al., 2010; Wu et al., 2016; Zhao et al., 2010; Zhou et al., 2010), we next monitored the expression of TCF1 in the various CD8 T cell subsets following YF-17D vaccination. First, by analyzing resting total CD8 T cells in a large number of donors (N=33), we observed a wide heterogeneity in TCF1 levels in human CCR7-CD8 T cell subsets (EM and EMRA). In line with mouse and human gene expression data (Crompton et al., 2015; Kratchmarov et al., 2018; Roychoudhuri et al., 2015; Willinger et al., 2006), we observed the hierarchical expression of TCF1: Naïve and memory subsets (including CM and SCM) expressed high levels of TCF1, while effector subsets (EM and EMRA) had low-to-negative levels of TCF1 (Figure 3). Similar to the inter-donor variability in subset distribution (Figure 3A and B), this single cell protein data in N=33 donors revealed that the fraction of TCF1+ cells was widely variable within CCR7-subsets across donors (EM and EMRA, Figure 3C and D).

**Figure 3.**
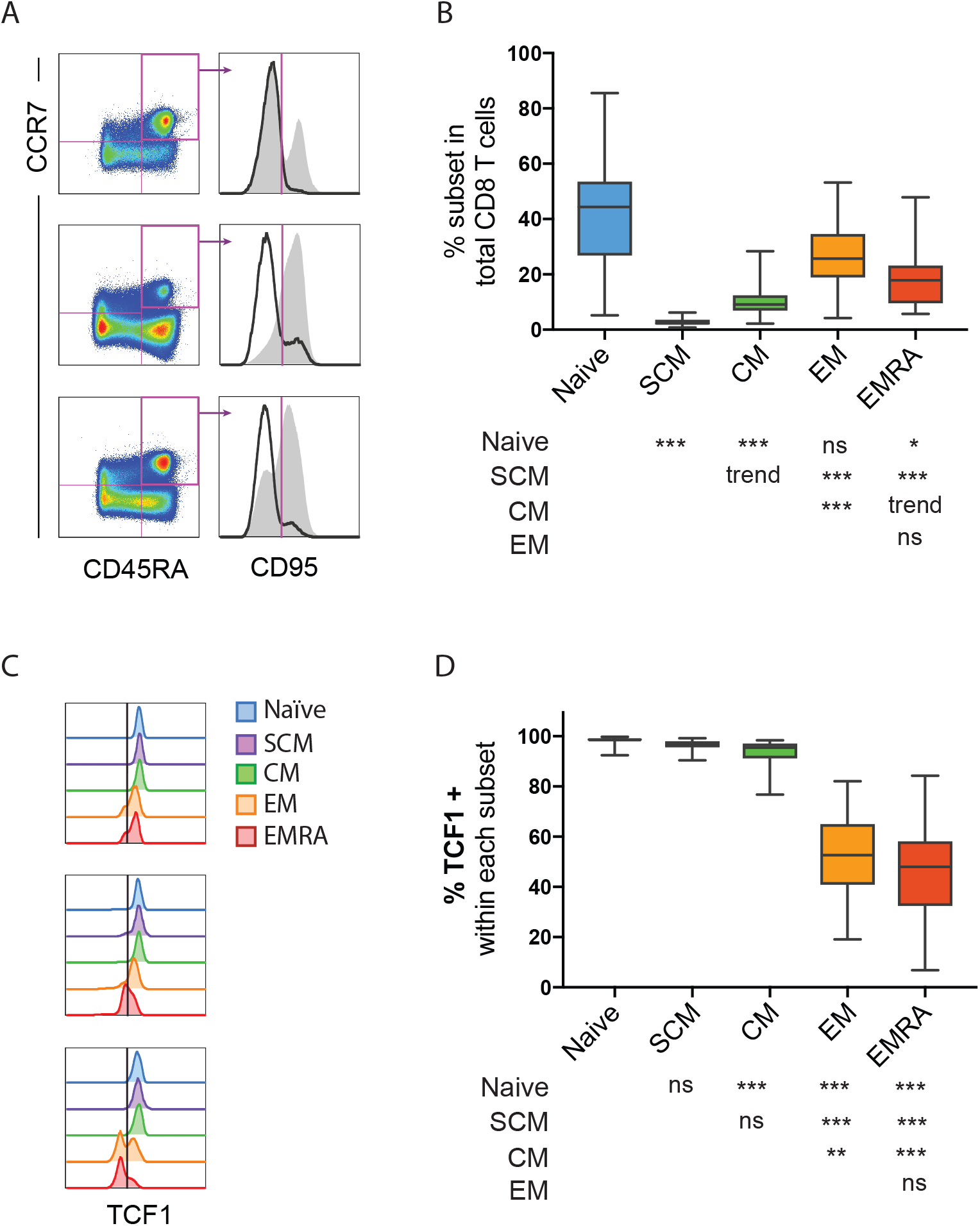
Patterns of subset distribution and TCF1 expression in human CD8 T cells across donors. **(A).** Flow cytometry analyses of CD8 T cells for subset composition showing 3 representative plots, and **(B)** quantitated subset frequencies in N=33 donors. **(C)**. Offset overlay histograms for TCF1 expression amongst CD8 T cell subsets showing 3 examples. (**D**) Frequencies for TCF1 expression in peripheral total CD8 T cell subsets from N=33 donors; these correspond to non-activated cells (unvaccinated or over 6 months after vaccination). Comparative p-values are shown in matrix format below each x-axis label, based on a Fridman test (non-parametric, paired): ns = not significant, trend = 0.05 to <0.10, * < 0.05, ** < 0.01, *** < 0.001.

We then analyzed TCF1 expression in A2/LLW-specific CD8 T cells at various time-points following vaccination with YF-17D. In order to assess the early and very long-term phases of the response, we analyzed samples from the longitudinal cohort (N=8, up to 6 months post-vaccination) together with samples from the cross-sectional cohort (N=26, up to 23.7 years post-vaccination, Table S1B: “Cross-sectional cohort: YF1 study” and Figure S3). In the first weeks, TCF1 positive frequencies sharply dropped in CCR7- subsets (EM and EMRA, Figure 4A and B), proving TCF1 downregulation during the acute response in humans in vivo. The maximum drop in TCF1 occurred at day 28 (Figure 4A and B), and appeared thus delayed relative to the activation peak at day 14 (Figure S2). After day 28, the CCR7- populations (EM or EMRA) showed a gradual increase in the percentages of TCF1+ cells, particularly visible in the decade-persisting EM and EMRA cells (Figure 4B). In contrast, we observed that TCF1 was maintained at high levels from baseline and throughout the observation time of two decades, in the three CCR7+ subsets: Naïve, SCM and CM (Figure 4A and B). Based on longitudinal modeling of the percentage of TCF1+ cells per subset and the comparison of the trends across subsets, the CCR7- subsets (EM and EMRA) were found to exhibit a distinct profile of TCF1 downregulation compared to TCF1 maintenance in CCR7+ subsets (Naïve, SCM, and CM) (Figure 4C). While CM cells showed a trend closer to the trends in Naïve and SCM, they were statistically distinct to all subsets; the trends of Naïve and SCM were not distinguishable (Figure 4C).

**Figure 4.**
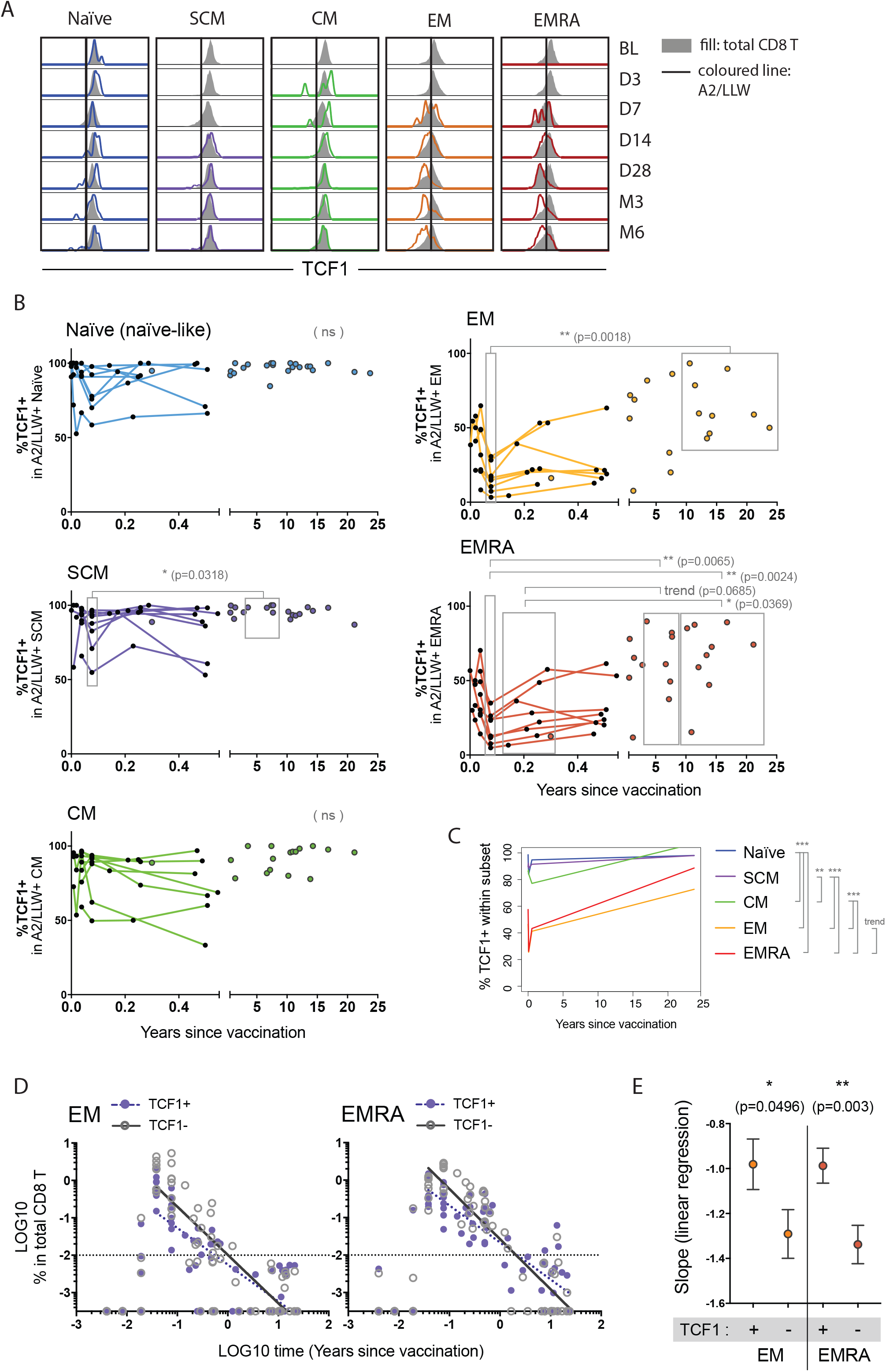
Dynamics of TCF1 expression in A2/LLW-specific CD8 T cell subsets in the early and late phases after YF-17D vaccination. **(A)** Flow cytometry profiles of TCF1 expression, longitudinally in the first 6 months after vaccination, in each subset. Overlay histograms show the A2/LLW multimer+ CD8 T cells in open line (colored per subset; absence denotes a non-detectable population) and the total CD8 T cell reference in grey fill below. Donor LAU 5081 is shown. **(B)** Frequencies of TCF1 expression in the various A2/LLW- specific CD8 T cell subsets, longitudinally in N=8 vaccinees in the early phase (first six months, line-connected dots per donor) and N= 26 vaccinees for the late phase (cross-sectional cohort: from 4 months to 23.7 years); total N=82 samples. P-values are based on Kruskal-Wallis (unpaired, non-parametric) for multiple comparisons amongst time-line groups, distributed as shown in Figure S4. **(C)** Statistical comparison of the % of TCF1+ cells per subset based on longitudinal modeling of the data (same dataset as in panel B) and Bonferroni adjustment of the pairwise p-values. **(D) and (E)** Frequencies of TCF1+ and TCF1- populations of A2/LLW-specific EM or EMRA subsets amongst total peripheral CD8 T cells. Corresponding linear regressions with least squares fit are shown for data from the peak of the response (at day 14). In D, the best-fit and standard error of the slopes from TCF1+ or TCF1- are compared, within each effector subset, with t-test p-values indicated. ns = not significant, trend = 0.05 to <0.10, * < 0.05, ** < 0.01, *** < 0.001.

Within the CCR7- subsets, we addressed whether the increase in the percentage of TCF1+ cells in the longer-term was linked to an overall or a relative increase in TCF1+ cells. We considered the frequencies of TCF1 positive or negative cells in each CCR7- subset in relation to the total CD8 T cells and from the peak of the response (day 14). We observed that: a) both TCF1+ and TCF1- populations declined with time (Figure 4D), and b) TCF1+ populations declined less than TCF1- populations (Figure 4D and E), in both EM and EMRA subsets. Rather than re-expression of TCF1 in CCR7- cells, these relative frequencies suggest that TCF1+ cells persist better than TCF1- cells in the long-term.

We further studied the expression of the Interleukin 7 Receptor alpha chain (IL7Ra) in the EMRA subset and found a pattern of IL7Ra expression globally correlated to TCF1 expression (Figure 5 and S5). In particular, both TCF1 and IL7Ra were co-enriched in EMRA cells persisting beyond six months and further co-enriched when persisting over three years. Similar trends were observed for the EM populations; however, because the EM subset in total CD8 T cells inherently features a substantial fraction of IL7Ra+ cells (as opposed to the scarcer fraction of IL7Ra+ in total EMRA), the TCF1 and IL7Ra co-enrichment was not significant; CCR7+ subsets express high levels of IL7Ra similar to high levels of TCF1+ (data not shown). This is in line with our previous analysis where A2/LLW-specific EMRA but not A2/LLW-specific EM showed significant enrichment of the IL7Ra as compared to their counterparts in total CD8 T cells (Fuertes Marraco et al., 2015).

**Figure 5.**
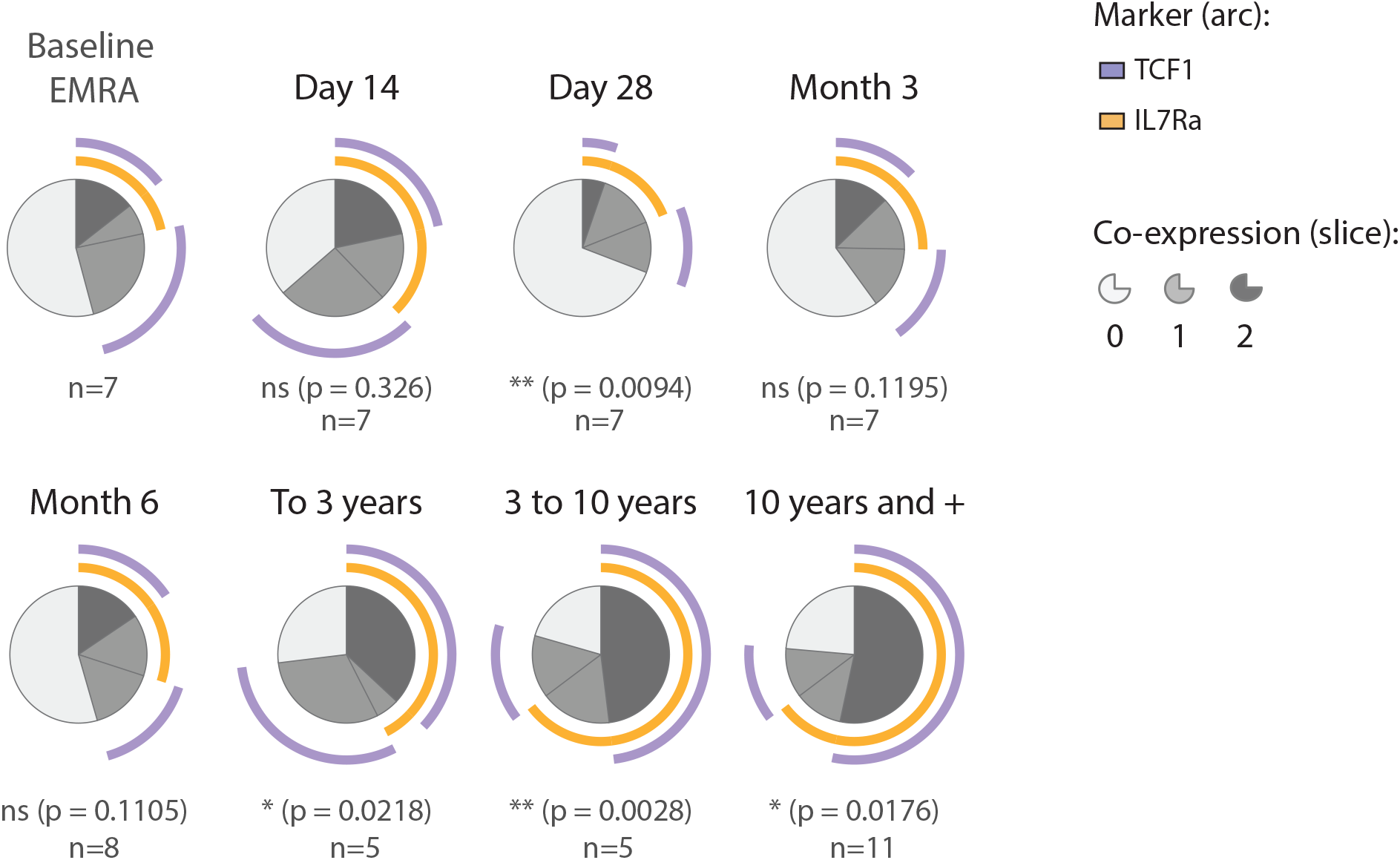
IL7Ra co-enriches with TCF1 in the long-term in A2/LLW-specific EMRA cells. Pie charts showing frequencies of the combinatorial expression of IL7Ra and TCF1 in the EMRA subset. Baseline EMRA correspond to EMRA from total CD8 T cells. Thereafter, post-vaccination EMRA populations that are A2/LLW-specific are shown (these are non- or insufficiently detectable for analysis before day 14). P-values (built-in t-test in SPICE, comparing to baseline): ns = not significant, * < 0.05, ** < 0.01, *** < 0.001.

### SCM CD8 T cells appear phenotypically close to the Naïve baseline, while effectors burst out of baseline and gradually contract

In order to detail the dynamics of the CD8 T cell response including multiple differentiation and activation markers, we applied multi-dimensionality reduction and unsupervised clustering to flow cytometry data using t-distributed Stochastic Neighbor Embedding (tSNE), and then further generated time-lapse representations. We applied this analysis strategy to samples from our longitudinal YF2 cohort, alone and in combination with long-term samples from the cross-sectional YF1 cohort. As detailed in the methods section, concatenated tSNE was possible for samples acquired with the same antibody panel and acquired under the same instrument configuration and normalized settings (Table S2).

First, tSNE was ran on single live total CD8 T cells from a pool of N=13 donors (non-acute samples), analyzing nine differentiation and activation markers: CCR7, CD45RA, CD95, TCF1, IL7Ra, PD1, CD69, CD38 and HLA-DR. The differentiation subsets were then gated using the standard strategy (Figure 6A, similar to Figure 1A) in order to locate them within the tSNE plots. The tSNE analysis of this CD8 T cell pool showed a distinct Naïve lobe, with SCM cells bridging this Naïve lobe into the remaining differentiation subsets, which were arranged in a gradient and formed a second lobe (Figure 6B: subset overlay, and C: individual subset populations). The localization of the subsets gated based on CCR7, CD45RA and CD95 (Figure 6A) corresponded well with the tSNE clustering, including the expression patterns expected for the remaining six markers (Figure 6D): for instance, IL7Ra and TCF1 were low while PD1 was high only in the CCR7- populations. In independent analyses, we analyzed N=16 donors, applying the 9-marker tSNE to each donor individually (Figure S6). We found that the pattern described above is reproducible across donors and tSNE runs, with donors showing variable sizes of the Naïve and differentiated lobes as expected based on the natural variability of the frequencies of CD8 T cell subsets (Figure 3). Of note, SCM where found in the bridge between lobes and also interspersed within the Naïve lobes.

**Figure 6.**
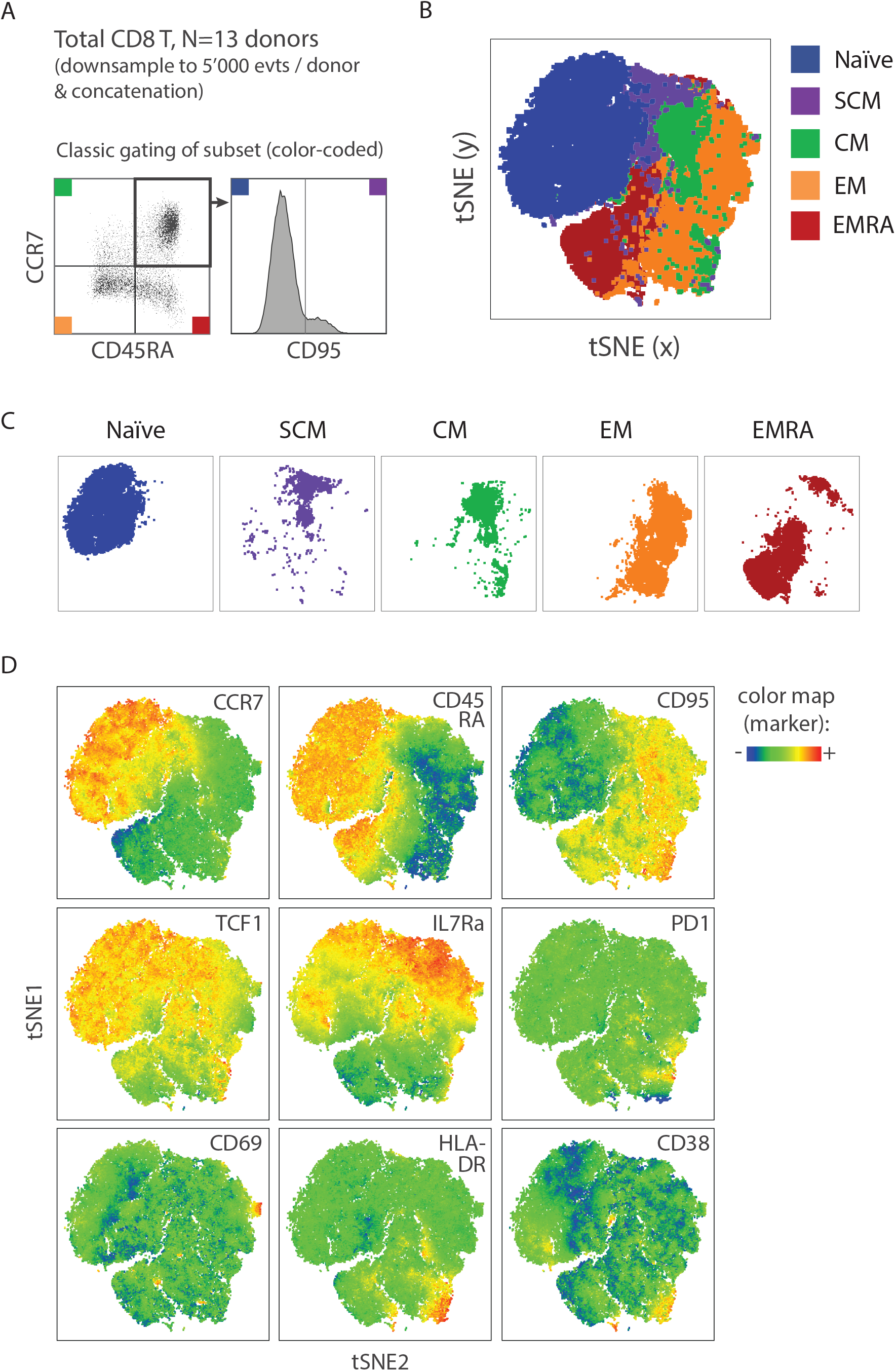
Gradient of differentiation in total CD8 T cells validated by t-Stochastic Neighbor Embedding (tSNE) of flow cytometry data. Total CD8 T cells from N=13 donors (ranging from 8.5 months to 23.7 years after vaccination, i.e. no acute phase samples) were analyzed with the tSNE plugin from FlowJo **(A)** Analysis strategy: single live CD8 T cells from each donor were downsampled to 5’000 events and the sum of N=13 donors were concatenated into a single file (70’000 events). This file was then gated and color-coded for the differentiation subsets as previously described (Figure 1A). **(B) and (C)** The N=13 concatenate was analyzed by tSNE using the plugin from FlowJo v10, reducing nine parameters (CCR7, CD45RA, CD95, TCF1, IL7Ra, PD1, CD69, HLA-DR, CD38) to two dimensions (tSNE x- and y-axes). Shown is the resulting unsupervised clustering tSNE plot, with the overlay (in B) or individual plots (in C) of the differentiation subsets gated as in panel A. **(D)** The tSNE plots showing the heatmap (based on median) of each marker, as indicated.

We next applied this 9-marker tSNE analysis to longitudinal series of YF-17D vaccination samples, running tSNE individually for each donor (N=7 longitudinal datasets). The subset overlay tSNE plots were generated as in Figure 6, by gating subsets in each time-point of the longitudinal concatenated file (Figure S7). We then represented the tSNE of each time-point and generated time-lapse animations of the data. We found a remarkable pattern of the dynamics of CD8 T cell differentiation during YF-17D vaccination across donors: SCM cells appear and remain very close to the location of baseline Naïve cells (Figure 7A; Video 1 showing N=7 subset overlays, and Video 2 showing each marker for donor LAU 5089 7 *video links inserted here*). In contrast, effector CCR7- populations burst out of the baseline Naïve location, peaking their distance at days 14-28, and gradually contracting.

**Figure 7.**
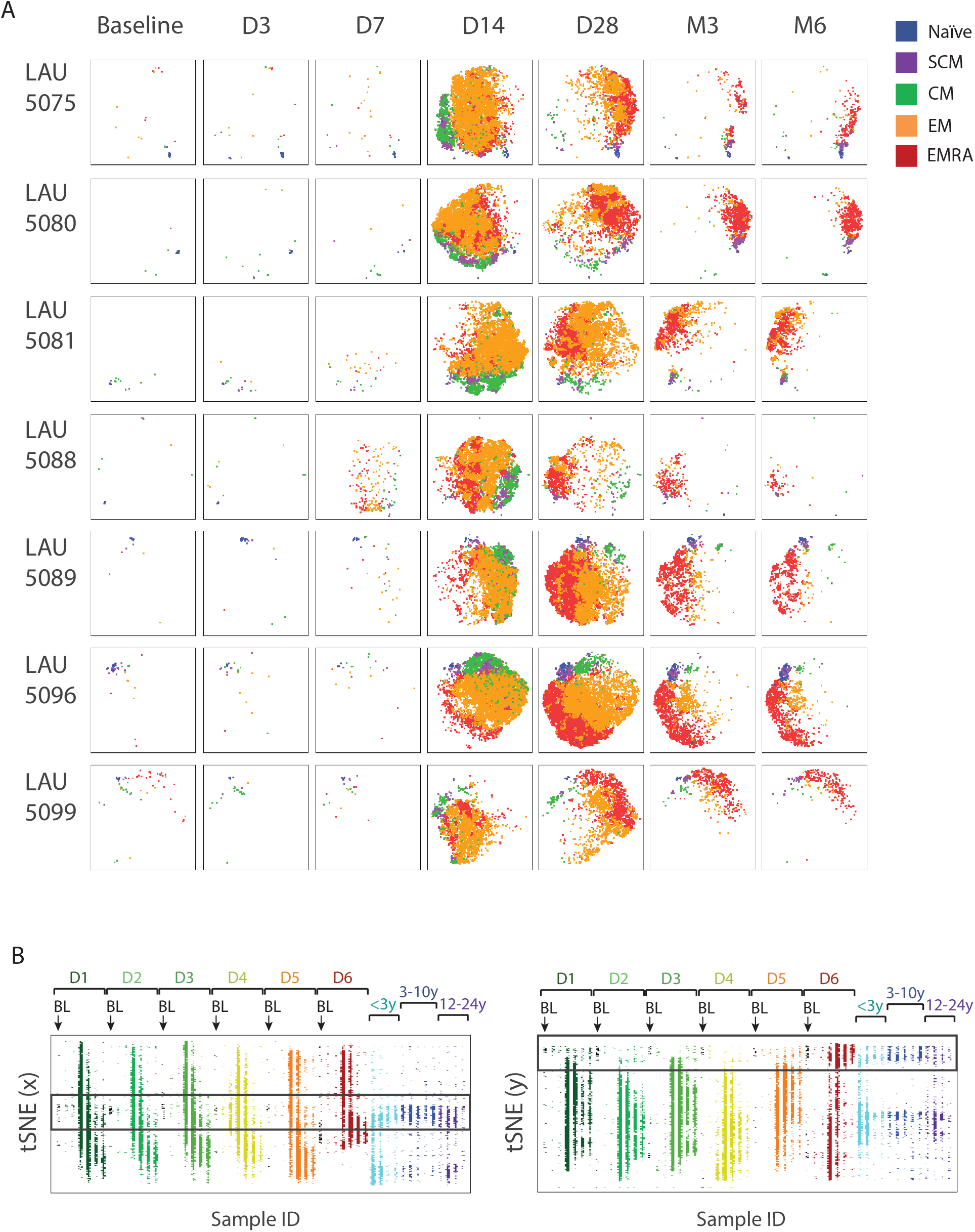
Time-lapse dynamics of CD8 T cell differentiation showing effector cell burst and permanent memory cell establishment during YF-17D vaccination. **(A)** N=7 donors were individually analyzed, performing tSNE analyses on the longitudinal data series: for each donor dataset, all the single live A2/LLW-specific CD8 T cell flow cytometry events acquired were concatenated and individually ran for tSNE. The resulting tSNE plots are shown for each time-point (gated based on sample ID). Samples from donor LAU 5096 were stained with a different antibody panel (« panel D ») compared to all other donors (stained with « panel C »), as detailed in Table S2. **(B)** Single live A2/LLW-specific CD8 T cells from a pool of N=55 samples acquired with the same flow cytometry panel and instrument configuration were concatenated and ran for tSNE. These included: N=6 donors (D1 to D6) with longitudinal data (7 time-points: BL, D3, D7, D14, D28, M3 and M6 in each sequence) together with N=13 donors from the cross-sectional cohort (grouped according to years since vaccination), as indicated. Shown are the plots of the calculated tSNE (either x- or y-dimension) versus sample ID. The black-bordered rectangle indicates the areas of permanency throughout vaccination.

To address how do decade-persisting CD8 T cells compare to the early dynamics, in further analyses, we concatenated N=6 longitudinal series of early vaccination samples (longitudinal cohort, up to 6 months; 7 time-points per series) with N=13 samples from the long-term, cross-sectional study. We found that in both tSNE dimensions (x and y), the long-term samples also featured a population that located in the baseline region, as a prolongation of the Naïve cells at baseline and the cells that retain phenotypic proximity to this baseline region throughout vaccination (Figure 7B).

## Discussion

To date, the differentiation pathways and fates of antigen-primed CD8 T cells are largely debated, a major question being whether memory cells establish in the first few days from precursors that diverge from effector fate or only later in the response from a fraction of acutely activated effector cells. With respect to the existing experimental evidence, one basic question still is: how early do memory subsets appear? Specifically concerning the more recently described SCM subset, the existing evidence is limited to one study using the macaque model of Simian Immunodeficiency Virus infection, where antigen-specific CD8 SCM cells are observed as early as day 7 of infection (supplementary data showing CM9/TL8-specific CD8 T cells in (Lugli et al., 2013)). Based on our clinical studies in YF-17D vaccinees, we show first evidence in humans, in vivo, that antigen-specific CD8 T memory cells including CM and SCM subsets are activated and expand during the acute phase of the response. The analysis of samples before day 14 was challenging due to the low frequencies of antigen-specific cells and due to the medical restrictions for blood withdrawals that precluded closer intervals in our study. Nevertheless, two donors showed rising memory cells as early as day 3 and 7, while virus was still detectable (day 3). Our data clearly exclude that memory subsets appear only once the antigen is cleared or after the acute peak, and provide evidence that cells with a memory phenotype establish early after priming, within the acute phase.

The particular value of this human experimental evidence is highlighted by the challenge in studying memory development based on phenotypic markers, and the fact that these markers globally vary between mouse and human systems. In the mouse, the major differentiation markers used are CD44 (for antigen-experienced) and CD62L (for Naïve and memory), as well as IL7Ra (naïve and memory including MPECs) and KLRG1 (terminally differentiated effectors and SLECs). Human CD8 T cell subsets are classically defined on the basis of CCR7 and CD45RA (as detailed in the introduction). The SCM subset was first identified in the mouse as cells within the classic naïve-like gate that distinctly express IL2Rb, Bcl-2, CXCR3, and SCA-1 (Gattinoni et al., 2009). Human SCM are distinguished from Naïve by the positive expression of CD95 and CD58, whereas these two markers are not used in mice. Conversely, the mouse SCA-1 has no human ortholog. The markers CXCR3, IL2Rb and Bcl-2 used in mice are also considered positive in human SCM (Fuertes Marraco et al., 2015; Gattinoni et al., 2011; Lugli et al., 2013; 2012). However, these three later markers are poorly discriminative to distinguish SCM from Naïve cells: i) Bcl-2 is highly expressed in both Naïve and SCM (Gattinoni et al., 2011) (in contrast to downregulation in cycling effectors cells (Miller et al., 2008)), ii) IL2Rb is higher in SCM but requires visualization with a second marker for Naïve/SCM discrimination, in contrast to the distinct CD95+ staining of SCM cells versus CD95- signal in naïve cells (Gattinoni et al., 2011), and iii) CXCR3 shows high inter-donor variability and substantial positive signal even in cells that are CD58 and CD95 negative such as Melan-A-specific CD8 T cells in healthy donors, which are predominantly naïve (Fuertes Marraco et al., 2015). A major challenge is thus the availability and choice of markers to distinctly define and visualize memory subsets. Ontogeny questions that require adoptive transfer and tracing is extremely limited in humans (only studies in the context of bone marrow transplants have successfully traced SCM generation from transferred T cells (Cieri et al., 2015)) and mouse models are largely used to study CD8 T cell differentiation. Notwithstanding, the discrepancy of differentiation markers used in different model systems makes human data uniquely informative, as observations are complementary but not fully transferrable across systems such as mouse and human.

Historically, it has been particularly challenging to distinguish SCM from Naïve cells: SCM cells represent a recently identified memory subset, hidden within the classic Naïve (“naïve-like”) gate. Interestingly, in our experiments, we found that there was a relatively constant level of antigen-specific CD8 T cells that fell in the naïve gate (CCR7- CD45RA- CD95-) even following priming. A hypothesis could be that these post-vaccination Naïve-gated cells have not actually been primed – this would require a compensatory replenishment of naïve cells with this antigen specificity, sufficiently rapid to immediately replenish the cells that have been primed and therefore depleted from the naïve pool. Murray et al calculated that after age 20, the naïve pool is maintained by homeostatic proliferation rather than thymic output (Murray et al., 2003) – all our donors were aged over 20, and it would be undistinguishable to know whether the naïve-gated cells proliferate due to homeostasis or due to priming. To our knowledge, in fact, there is to date no marker that can definitely prove that a given CD8 T cell has been primed. In the mouse, CD44 is often used as a marker to distinguish differentiated cells, yet there is no proof that CD44 expression is truly correlative of antigen priming experience; there is neither no such equivalent marker conventionally used in human experiments. In previous studies including ours, there is evidence mainly substantiated from whole transcriptomic profiles, epigenetic imprints and functional assays, that subsets are arranged in a gradient, ordered from Naïve to SCM, CM, EM and EMRA (Crompton et al., 2015; Fuertes Marraco et al., 2015; Gattinoni et al., 2011; Mahnke et al., 2013). Intriguingly, we did observe that a substantial portion of cells in the naïve gate were undergoing activation, clearly visible at the peak of the response. In line with the argument that SCM amongst antigen-experienced cells preserve highest “naïveness”, we hypothesize that the cells that remain in the naïve gate after priming may have been effectively primed but are memory cells that preserve a phenotype that is very close to the naïve, even closer than SCM. Recently, Costa del Amo et al. found subpopulations of SCM cells with distinct turn-over rates in vivo (Costa del Amo et al., 2018), which highlights further potential heterogeneity within subset gates, and in support of the differentiation continuum from naïve to memory to effector.

In our animated longitudinal tSNE analyses on nine standard activation and differentiation markers, it was particularly visible that a fraction of cells remained in the region where cells (Naïve) located at baseline. We observed YF-specific CD8 T cell subsets with phenotypes of naïve, memory (CM and SCM, CCR7+) or effector / effector memory (EM or EMRA, CCR7+) cells, and each of these subsets showed activation at the peak of the response and downregulated activation markers at later time-points. This observation highlights the importance of distinguishing between displaying a memory or effector phenotype and being in an activated or resting state. The progressive differentiation model does account for activated / effector phenotypes that may rise from each of the subsets (Mahnke et al., 2013). To-date, methodologies used in the study of CD8 T cell differentiation include the definition of memory cells solely on the basis of the time of sampling (meaning that all cells that are detected after the acute phase are memory cells), including studies in humans vaccinated with YF-17D (Akondy et al., 2017). In fact, our analyses show that there is wide heterogeneity in differentiation phenotypes very early on, with SCM cells appearing as a population that is phenotypically close to the naïve baseline and distinct from the burst of effector subsets at the peak of the response. Along these lines, a study in mice showed that CD8 T cells that have undergone the first division upon priming *in vivo* display transcriptional heterogeneity, with two main clusters with effector-like or memory-like profiles(Kakaradov et al., 2017). Our data shows phenotypic heterogeneity also in the very long-term, evidenced by the fact that EMRA cells are detectable decades after vaccination as a fraction of cells separate from SCM cells (Figure 1, Figure S3) (Fuertes Marraco et al., 2015). Even though they are phenotypically quite distinct (SCM versus EMRA), stem cell features such as long-term persistence and self-renewal are likely shared in these long-term populations, at least in a fraction of them. Interestingly, we found that it is the TCF1-expressing cells that preferentially persisted in the range of years-to-decades. This was pertinent not only for the SCM subset (TCF1 high from baseline and permanently thereafter) but also particularly visible in the fraction of TCF1+ cells within the EMRA subset that preferentially persisted long-term over TCF1- EMRA. The latter suggests that TCF1 may generally support cellular persistence and thus also the maintenance of long-term effector cells that are readily available in the event of reinfection.

Another historical challenge is that the classic nomenclature of differentiation subsets based on CCR7 and CD45RA was primarily defined studying resting human T cells, where all non-naïve subsets are termed “memory”. This nomenclature does not phenotypically distinguish acutely activated effectors (CCR7-) from the “memory”-termed effector subsets EM (CCR7- CD45RA-) and EMRA (CCR7- CD45RA+) (Mahnke et al., 2013). Activation and cycling markers may distinguish acute phase effectors versus resting / long-term EM and EMRA: activated CCR7- cells (HLA-DR+ CD38+) would be effectors, and CCR7- cells that are HLA-DR- and CD38- would be EM/EMRA. However, how do we define the cells that show a combined memory (CCR7+) and activated phenotype, such as the SCM and CM subsets that we detected in the acute phase being as activated as effector CCR7- subsets? The longitudinal phenotyping we hereby present builds on the current nomenclature and marker definition of human CD8 T cell subsets and extends it by considering the activated (acute phase) versus resting states in complement with memory and/or effector markers.

Altogether, based on clinical studies on YF-17D vaccination, we provide first evidence in humans, *in vivo*, on the early appearance of SCM CD8 T cells. The SCM phenotype that predominantly and stably persists in the decade long-term is detectable within the first week, and shows activation and expansion during the early acute phase. The results support differentiation models where memory cells arise very early without an obligatory transition through a full effector phenotype stage, yet showing an activated state on top of a memory phenotype. This would be in line with the existence of a continuum of differentiation phenotypes, where long-term memory cells diverge from the full-blown effector burst and persist by preserving highest “naïveness” (proximity to the Naïve).

## Acknowledgements

We thank Benton Lawson and the Centre for AIDS Research of Emory University (US) for the quantification of YFV load in plasma samples and the Flow Cytometry Facility of the University of Lausanne (CH) for cytometer instrument configuration and maintenance. We thank Nicole Montandon for technical assistant in processing blood specimens, and Paula Marcos Mondéjar for participating in the longitudinal study coordination. We thank Blaise Genton, Francine Widmer, Pierrette Meige and the personnel of the ‘Centre de vaccination et médecine des voyages’ at the PMU for the coordinated efforts and collaboration with us to receive the donors and withdraw blood specimens for the longitudinal study. Finally, we warmly thank all donors that volunteered and thus preciously contributed to our findings. This study was funded by Ludwig Cancer Research and the Cancer Research Institute (both N.Y., U.S.A), the Swiss National Science Foundation (grants: 320030_152856 to DES, 310030-179459 to DES, 310030B_179570/B to WH).

## Author Contributions Statement

SAFM, AB, WH and DES conceived and designed experiments, SAFM, HMEH, HOS, and DES elaborated the clinical study protocols, SAFM and AB performed experiments, SAFM, AB and SN analyzed data. All authors revised and approved the final version of the manuscript.

---

**Video 1. “N=7 subsets”: Dynamics of the differentiation of A2/LLW-specific CD8 T cells during YF-17D vaccination, showing subset composition for N=7 donors.** Time-lapse animation of the longitudinal tSNE analysis with subset overlays in N=7 donors, as indicated. Each donor sequence is spaced by 1 second, showing subset composition, and starting from Baseline, then Day 3, Day 7, Day 14, Day 28, circa 3 Months and circa 6 Months after YF-17D vaccination. For each donor, all the single live A2/LLW-specific CD8 T cell events acquired were concatenated and ran for the 9-marker tSNE.

**Video 2. “LAU 5089 markers”: Dynamics of the differentiation of A2/LLW-specific CD8 T cells during YF-17D vaccination, showing each of the 9 markers for vaccinee LAU 5089.** Time-lapse animation of the longitudinal tSNE analysis of donor LAU5089, showing sequences starting from Baseline, then Day 3, Day 7, Day 14, Day 28, Day 84 (ca. 3 months) and Day 185 (ca. 6 months) after YF-17D vaccination. Each sequence is spaced by 1 second, and shows subset overlay or the indicated heatmapped marker. All the single live A2/LLW-specific CD8 T cell events acquired were concatenated and ran for the 9-marker tSNE.

## Supplementary Material

**Figure S1. Longitudinal differentiation of A2/LLW-specific CD8 T cells in donors with detectable Yellow Fever viral load.** Data are quantified as in Figure 1D, showed for each individual donor that showed positive Yellow Fever viral load (N=5 out of 8 donors).

**Figure S2. Longitudinal analysis of activation markers in A2/LLW-specific CD8 T cell subsets.** The analysis is performed as in Figure 2, showing all time-points. The pie charts are translucent for the time-points and subsets with less than 3 donors with interpretable data. N values are indicated below each pie chart. P-values (built-in t-test in SPICE, comparing subsets within each time-point): ns = not significant, * < 0.05, ** < 0.01, *** < 0.001.

**Figure S3. Frequencies of A2/LLW-specific CD8 T cell subsets in subjects of the cross sectional cohort.** The frequency of A2/LLW-specific CD8 T cells within each subset is shown for the cross-sectional donors (analyzed in Figures 3 to 7 in combination with donors of the longitudinal cohort).

**Figure S4: Expression of TCF1 in A2/LLW-specific CD8 T cells of donors from the longitudinal and cross-sectional cohorts.** Donors were grouped according to various time intervals since vaccination, each dot representing one donor. P-values: ns = not significant, trend = 0.05 to <0.10, * < 0.05, ** < 0.01, *** < 0.001, after Kruskal-Wallis multiple comparisons (unpaired, non-parametric).

**Figure S5. TCF1 and IL7Ra co-expression in total and A2/LLW-specific EMRA cells early and long-term after YF-17D vaccination.** Data are analyzed as in Figure 5, showing the data corresponding to either total or A2/LLW-specific EMRA cells, in the various time-point groups. P-values (built-in t-test in SPICE, comparing total vesus A2/LLW-specific within each time-point group): ns = not significant, trend = 0.05 to <0.10, * < 0.05, ** < 0.01, *** < 0.001.

**Figure S6. Individual yet conserved tSNE pattern of differentiation subsets in total CD8 T cells.** A downsample of 75’000 single live total CD8 T cells was exported for each of the N=16 donors, individually running tSNE and analyzing each donor (gated and represented as in Figure 6B). The first N=13 donors correspond to the group analyzed in Figure 6 (ranging from 8.5 months to 23.7 years after vaccination, i.e. no acute phase samples); data of N=3 additional donors (ranging from 10.5 to 13.8 years after vaccination) originate from a different antibody panel configuration (“panel D”, see Table S2).

**Figure S7. Longitudinal dynamics of subset composition and marker expression from the tSNE analysis of donor LAU 5089. (A)** For each donor dataset, all the single live A2/LLW851 specific CD8 T cell flow cytometry events were concatenated and individually ran for tSNE: this generates a tSNE plot of all events. Each time-point is then gated based on sample ID, and subsets further gated and color-coded as detailed in Figure 6A. **(B)** Longitudinal tSNE plots showing the heatmap (based on median) of each marker, as indicated.

## Supplementary figure legends

Table S1

Table S2

Table S3

