## Supplemental Figures for "The human CD8 T stem cell-like memory phenotype appears in the acute phase in Yellow Fever virus vaccination"

Figure S1

LAU 5080

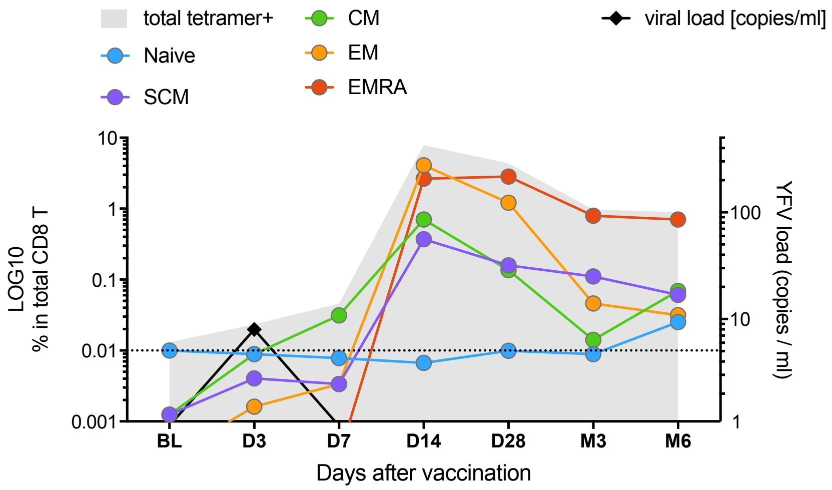


LAU 5081

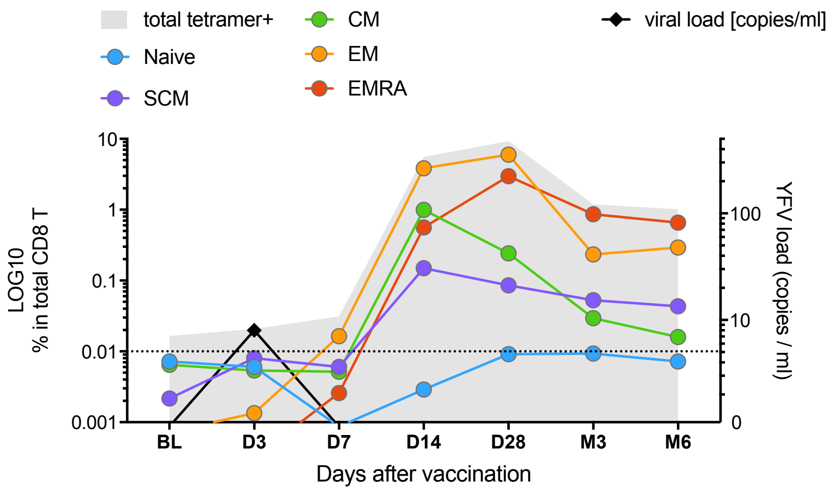


LAU 5088

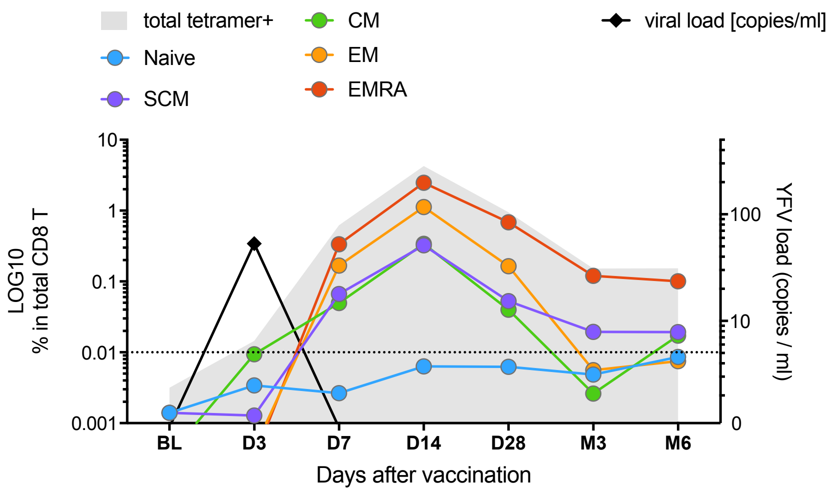

### Figure S1 (contd)

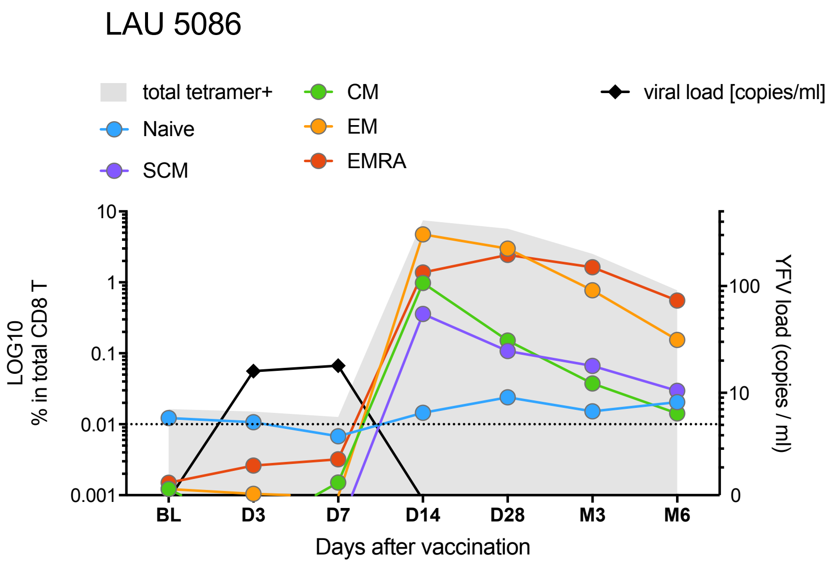


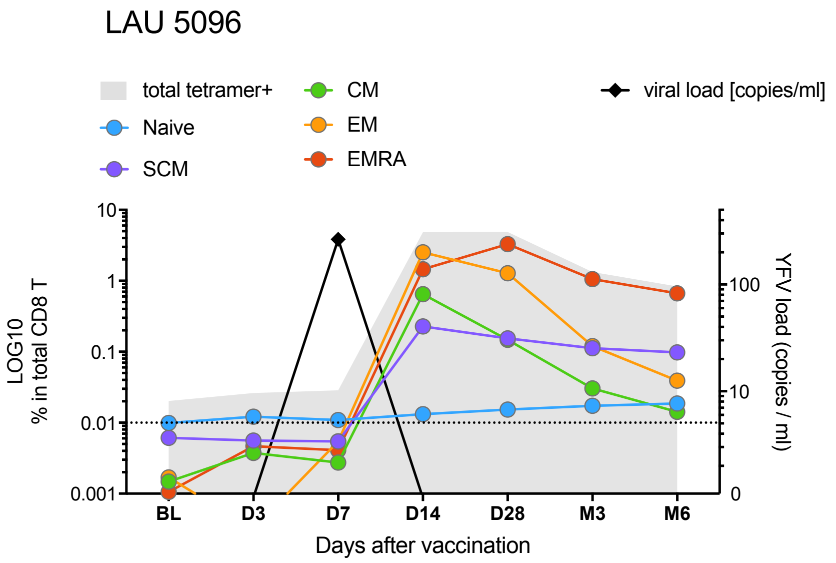

Figure S2

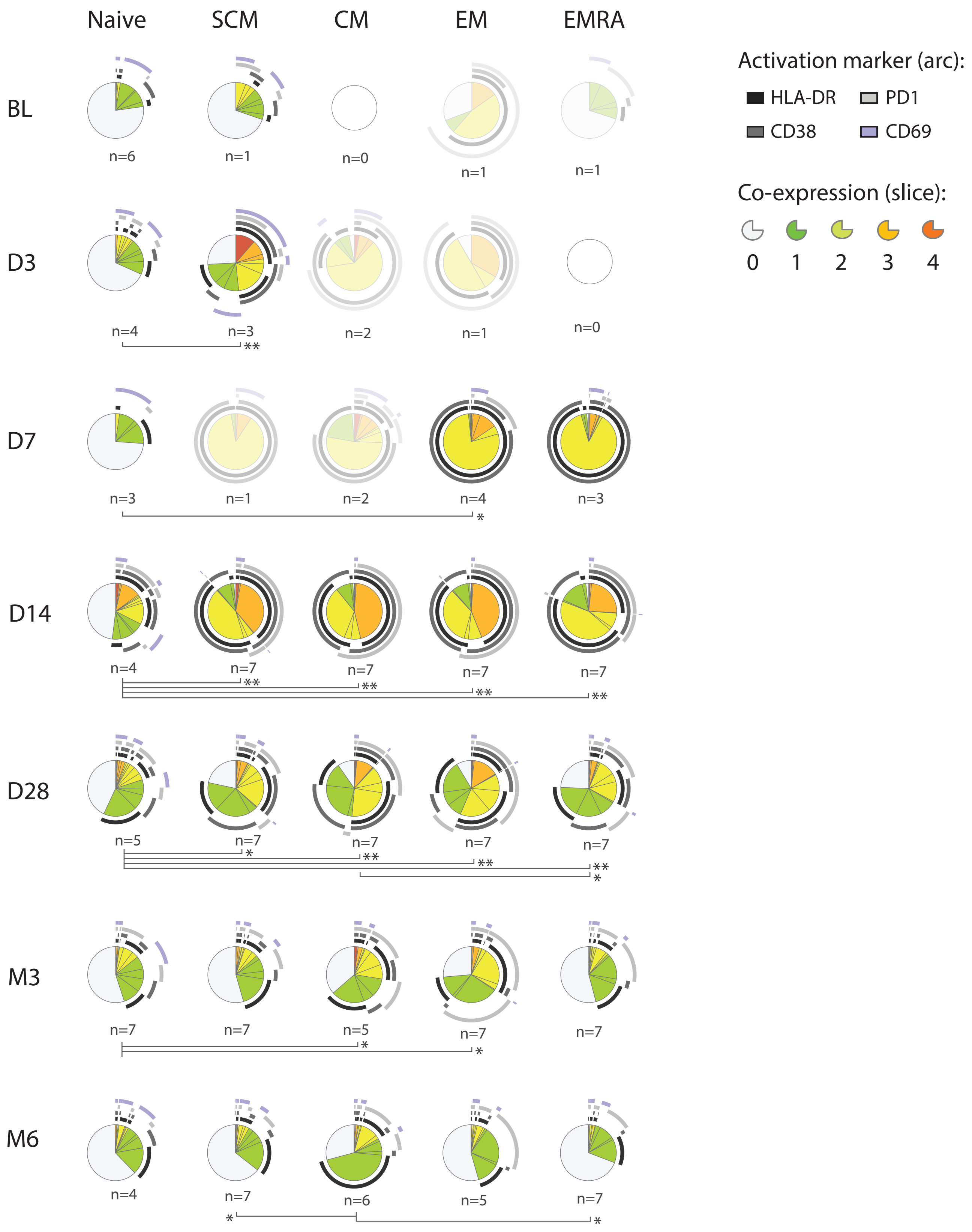

Naive
SCM
CM
EM
EMRA
BL
D3
D7
D14
D28
M3
M6
n=6
n=1
n=0
n=1
n=1
n=4
n=3
n=2
n=1
n=0
n=3
n=1
n=2
n=4
n=3
n=4
n=7
n=7
n=7
n=7
n=5
n=7
n=7
n=7
n=7
n=7
n=7
n=5
n=7
n=7
n=4
n=7
n=6
n=5
n=7
Activation marker (arc):
HLA-DR
PD1
CD38
CD69
Co-expression (slice):
0
1
2
3
4

Figure S3

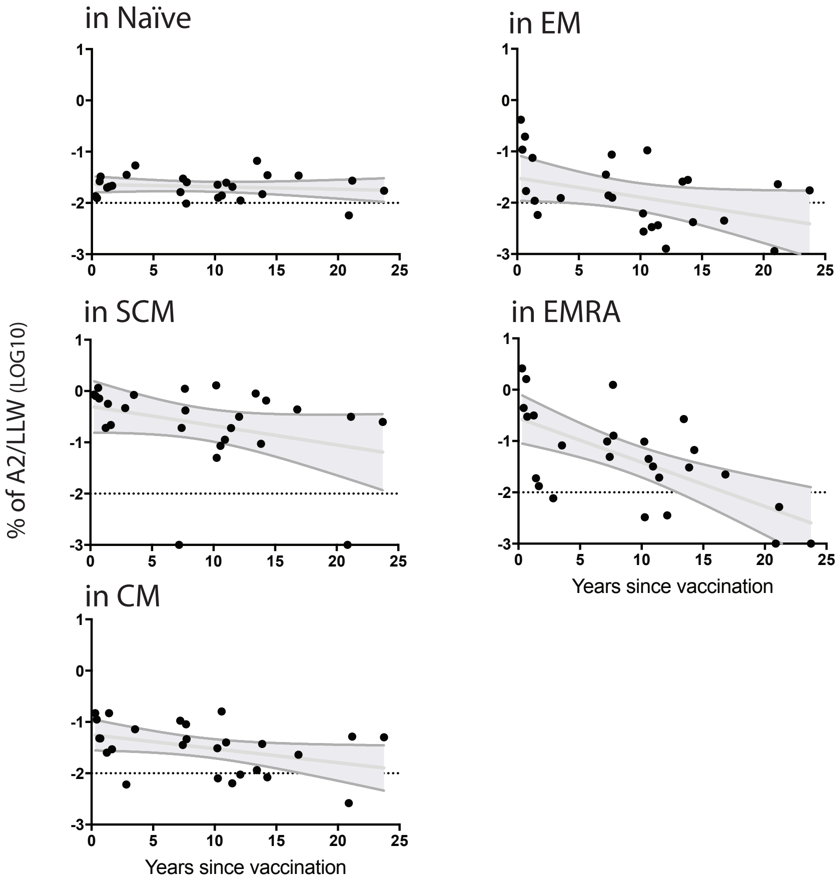

in Naïve
in EM
in SCM
in EMRA
in CM
% of A2/LLW (LOG10)
Years since vaccination
Years since vaccination
1
0
-1
-2
-3
0
5
10
15
20
25

Figure S4

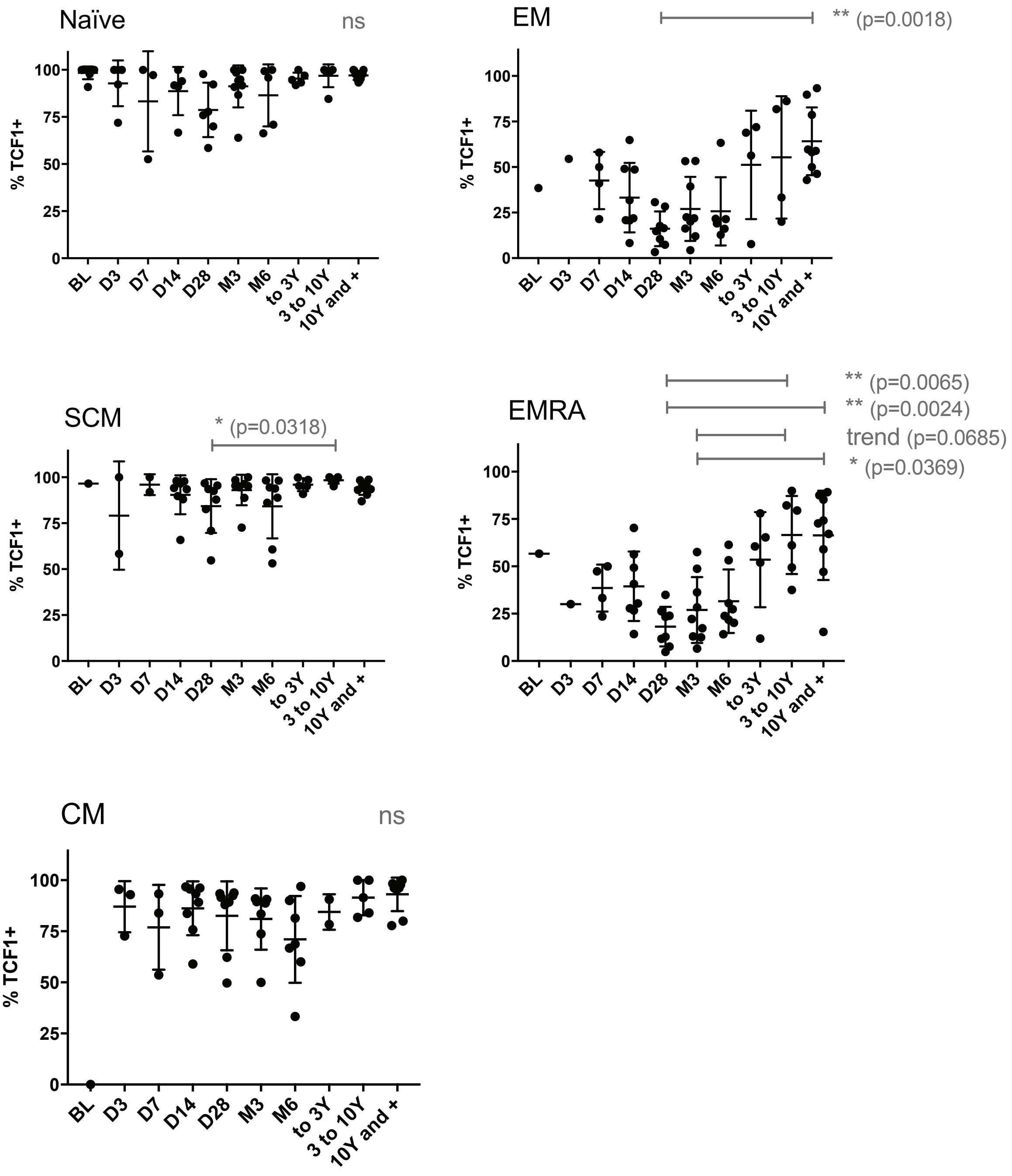
Naïve
ns
EM
** (p=0.0018)
SCM
* (p=0.0318)
EMRA
** (p=0.0065)
** (p=0.0024)
trend (p=0.0685)
* (p=0.0369)
CM
ns
% TCF1+
0
25
50
75
100
BL
D3
D7
D14
D28
M3
M6
to 3Y
3 to 10Y
10Y and +

Figure S5

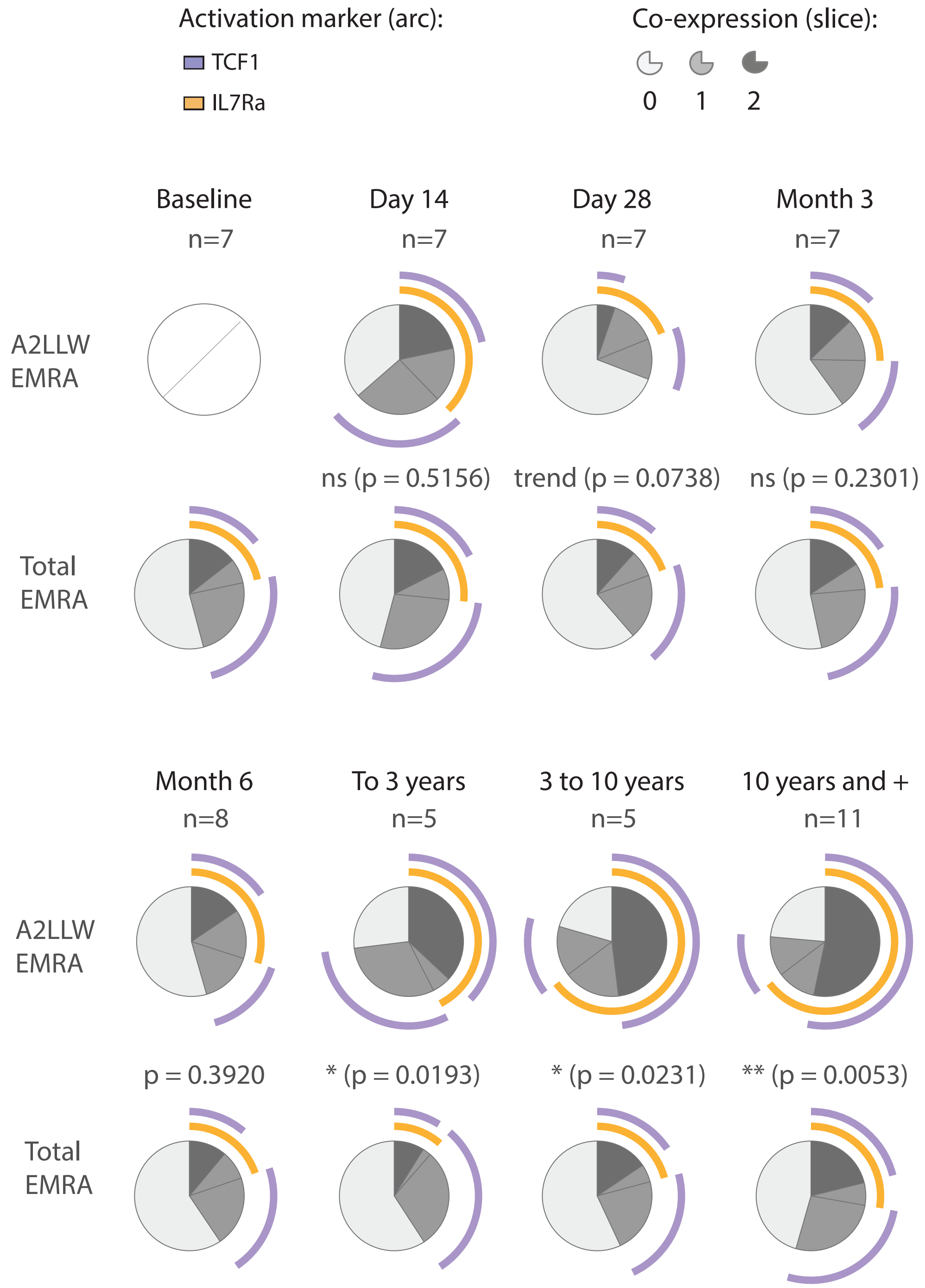
Activation marker (arc):
TCF1
IL7Ra
Co-expression (slice):
0
1
2
Baseline
n=7
Day 14
n=7
Day 28
n=7
Month 3
n=7
A2LLW
EMRA
ns (p = 0.5156)
trend (p = 0.0738)
ns (p = 0.2301)
Total
EMRA
Month 6
n=8
To 3 years
n=5
3 to 10 years
n=5
10 years and +
n=11
A2LLW
EMRA
p = 0.3920
* (p = 0.0193)
* (p = 0.0231)
** (p = 0.0053)
Total
EMRA

Figure S6

Total CD8 T, n=16 donors (n=13 from panel 1, n=3 from panel 2)
(downsample to 75'000 evts / donor)

with classic gating for color-coded subset overlay

each donor ran individually
(tSNE run per donor)

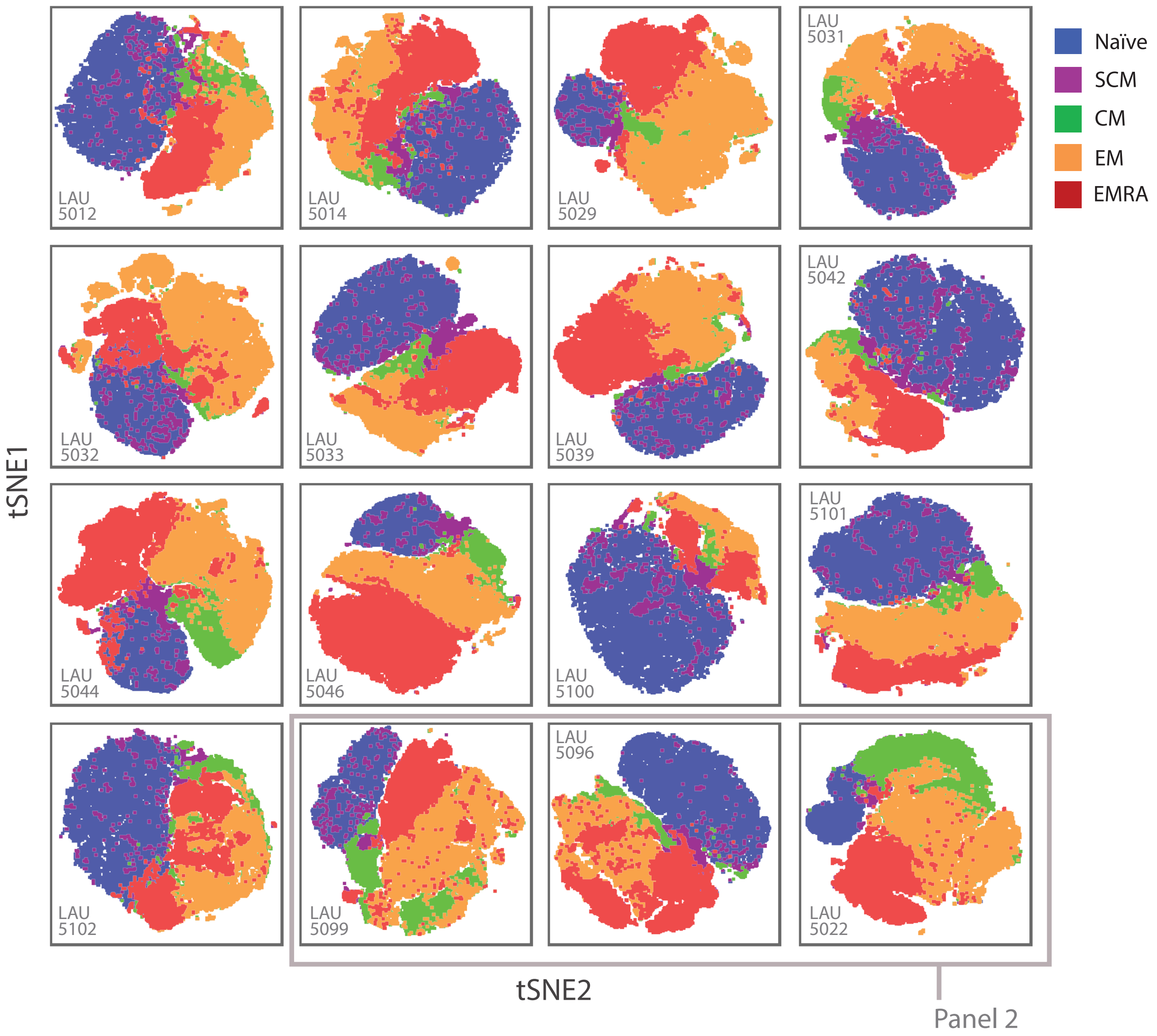

Figure S7 + animation links LAU 5089

A

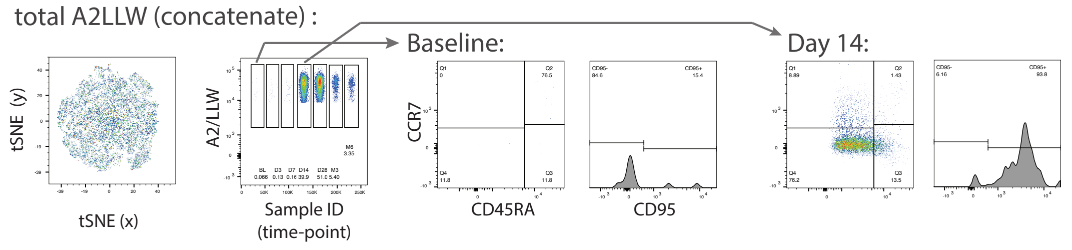


B

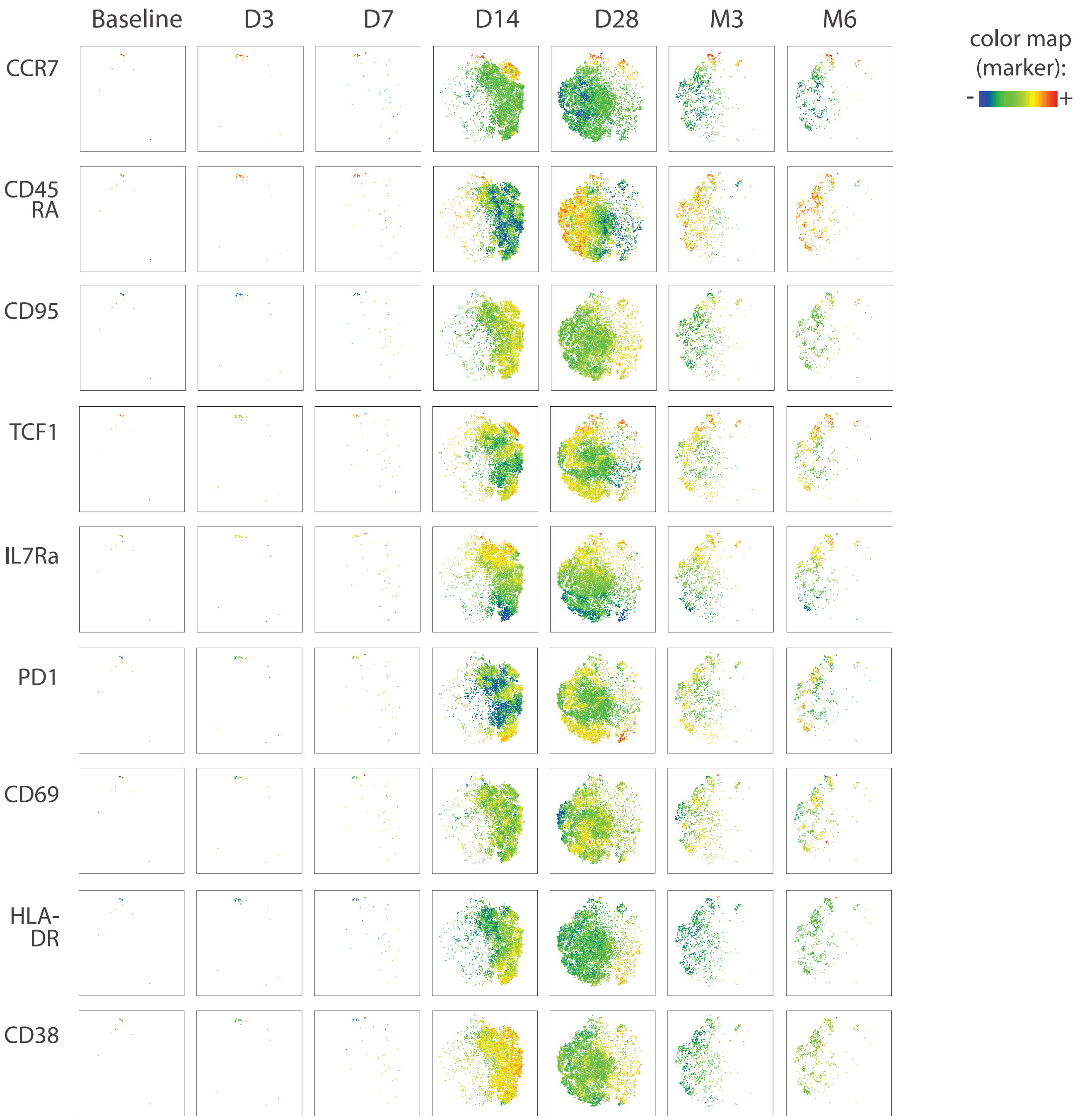
