## Supplemental Tables for "The human CD8 T stem cell-like memory phenotype appears in the acute phase in Yellow Fever virus vaccination"

Table S1. Metadata: donors analysed in this study

### A. Longitudinal Cohort: "YF2" study

| Donor | Age | Gender | Vaccine history<br>*Days* |
|---|---|---|---|
| **LAU5075** | 22.74 | female | -46 |
| | 22.87 | | 3 |
| | 22.89 | | 7 |
| | 22.90 | | 14 |
| | 22.94 | | 28 |
| | 23.04 | | 63 |
| | 23.36 | | 182 |
| **LAU5080** | 47.43 | female | -21 |
| | 47.50 | | 3 |
| | 47.51 | | 7 |
| | 47.53 | | 14 |
| | 47.57 | | 28 |
| | 47.75 | | 94 |
| | 48.00 | | 185 |
| **LAU5081** | 39.76 | female | -39 |
| | 39.88 | | 3 |
| | 39.89 | | 7 |
| | 39.91 | | 14 |
| | 39.95 | | 28 |
| | 40.13 | | 94 |
| | 40.36 | | 178 |
| **LAU5086** | 23.10 | male | -21 |
| | 23.16 | | 3 |
| | 23.18 | | 7 |
| | 23.20 | | 14 |
| | 23.23 | | 28 |
| | 23.30 | | 52 |
| | 23.62 | | 168 |
| **LAU5088** | 22.98 | male | -21 |
| | 23.05 | | 3 |
| | 23.06 | | 7 |
| | 23.08 | | 14 |
| | 23.12 | | 28 |
| | 23.33 | | 105 |
| | 23.58 | | 199 |
| **LAU5089** | 33.49 | female | -21 |
| | 33.55 | | 3 |
| | 33.56 | | 7 |
| | 33.58 | | 14 |
| | 33.62 | | 28 |
| | 33.77 | | 84 |
| | 34.05 | | 185 |
| **LAU5096** | 21.59 | male | -25 |
| | 21.66 | | 3 |
| | 21.67 | | 7 |
| | 21.69 | | 14 |
| | 21.73 | | 28 |
| | 21.86 | | 77 |
| | 22.12 | | 171 |
| **LAU5099** | 34.59 | male | -35 |
| | 34.69 | | 3 |
| | 34.70 | | 7 |
| | 34.72 | | 14 |
| | 34.76 | | 28 |
| | 34.93 | | 91 |
| | 35.18 | | 182 |

Longitudinal sampling:
- in order to meet the limitations of blood volume donation per month, the baseline sample was collected at least 21 days before vaccination
- the targeted post-vaccination time-point ranges were: day 3 +/-0, day 7 +/-0, day 14 +/-1, day 28 +/-2, 3 months +/- 2 weeks, 6 months +/- 2 weeks

(Table S1 continued)

**B. Cross-sectional cohort: "YF1" study (Fuertes Marraco et al. 2015)**

| | | | Vaccine history |
|---|---|---|---|
| **Donor** | Age | Gender | *Years* |
| **LAU 5051** | 35.60 | female | 0.30 |
| **LAU 5013** | 29.6 | male | 0.43 |
| **LAU 5019** | 31.00 | female | 0.63 |
| **LAU 5044** | 51.40 | female | 0.71 |
| **LAU 5013 (leukapheresis)** | 30.4 | male | 1.24 |
| **LAU 5046** | 60.80 | male | 1.43 |
| **LAU 5031** | 24.70 | female | 1.66 |
| **LAU 5039** | 24.40 | male | 2.82 |
| **LAU 5035** | 22.5 | female | 3.53 |
| **LAU 5042** | 24.0 | female | 6.67 |
| **LAU 5001 (leukapheresis)** | 26.8 | female | 7.20 |
| **LAU 5048** | 33.1 | male | 7.41 |
| **LAU 5012** | 29.5 | female | 7.68 |
| **LAU 5058** | 29.8 | female | 7.71 |
| **LAU 5014** | 21.6 | female | 10.21 |
| **LAU 5036** | 21.0 | female | 10.26 |
| **LAU 5100** | 32.4 | female | 10.56 |
| **LAU 5057** | 26.9 | female | 11.05 |
| **LAU 5033** | 22.9 | female | 11.43 |
| **LAU 5022** | 40.8 | male | 12.08 |
| **LAU 5043** | 70.1 | male | 13.42 |
| **LAU 5102** | 25.3 | female | 13.85 |
| **LAU 5032** | 51.1 | male | 14.27 |
| **LAU 5029** | 25.8 | male | 16.80 |
| **LAU 5002** | 42.4 | male | 20.88 |
| **LAU 5028** | 61.4 | female | 21.16 |
| **LAU 5101** | 24.6 | female | 23.74 |

Table S2. Flow Cytometry antibody panels and distribution of analysis of donors

### A. Flow cytometry panels used in this study

Cytometer Gallios :

| Fluorochrome | Panel **A** | Panel **B** |
|---|---|---|
| FITC | CD58 | PD1 |
| PE | iTCF1 (aRab) | iTCF1 (aRab) |
| ECD | CD45RA | HLA-DR |
| PerCP-Cy5.5 | - | CD127 |
| PerCP-eF710 | PD1 | - |
| PC7 | CD95 | CD95 |
| APC | A2/LLW | A2/LLW |
| A700 | CD28 | CD45RA |
| APC-A750 / nearIR | CD8 | - |
| Near InfraRed | - | Vivid nIR |
| BV421 | CCR7 | CCR7 |
| AmCyan | Vivid Aqua | - |
| Krome Orange | CD16 KrO | CD8 |

Cytometer LSR-II SORP :

| Fluorochrome | Panel **C** | Panel **D** |
|---|---|---|
| FITC | iTCF1 (aRab) | iTCF1 (aRab) |
| PerCP-Cy5.5 | - | CD127 |
| PerCP-eF710 | PD1 | - |
| PE | A2/LLW | A2/LLW |
| ECD | HLA-DR | CD45RA |
| PE-Cy7 | CD95 | CD95 |
| APC | CD58 | HLA-DR |
| A700 | CD38 | CD38 |
| APC-A750 | CD8 | CD8 |
| BrV421 | CCR7 | CCR7 |
| BV510 | CD45RA | - |
| BV605 | CXCR5 | CXCR5 |
| BV650 | CD69 | CD69 |
| BV711 | - | PD1 |
| BV785 | CD127 | - |
| eF455-UV | Fixable viability Dye | Fixable viability Dye |

i = intracellular
aRab = secondary anti-rabbit

(Table S2 continued)

### B. Distribution of donors across flow cytometry panels

| Donor | Exp | Time since vaccination | Panel |
|---|---|---|---|
| LAU 5051 | SF-2014-09-19 | 0.3 | **A** (Gallios) |
| LEUKA 5013 | | 1.24 | |
| LAU 5001 (Leukapheresis) | | 7.2 | |
| LAU 5057 | SF-2015-05-12 | longitudinal | **B** (Gallios) |
| LAU 5086 | | | |
| LAU 5013 (std) | SF-2017-03-30 | 0.43 | **C** (LSRII SORP, 2016) |
| LAU 5019 | | 0.63 | |
| LAU 5035 | | 3.53 | |
| LAU 5048 | | 7.41 | |
| LAU 5058 | | 7.71 | |
| LAU 5036 | | 10.26 | |
| LAU 5002 | | 20.88 | |
| LAU 5028 | | 21.16 | |
| LAU 5075 | AB-2017-06-02 | longitudinal | **C** (LSRII SORP, May 2017) |
| LAU 5089 | | | |
| LAU 5081 | AB-2017-06-08 | longitudinal | |
| LAU 5099 | | | |
| LAU 5080 | AB-2017-06-14 | longitudinal | |
| LAU 5088 | | | |
| LAU 5044 | SF-2017-09-27 | 0.71 | **C** (LSRII SORP, Aug2017) |
| LAU 5046 | | 1.43 | |
| LAU 5031 | | 1.66 | |
| LAU 5039 | | 2.82 | |
| LAU 5042 | | 6.67 | |
| LAU 5012 | | 7.68 | |
| LAU 5014 | | 10.21 | |
| LAU 5033 | | 11.43 | |
| LAU 5043 | | 13.42 | |
| LAU 5032 | | 14.27 | |
| LAU 5029 | | 16.80 | |
| LAU 5100 | AB-2015-09-27 | 10.56 | |
| LAU 5101 | | 23.74 | |
| LAU 5102 | | 13.85 | |
| LAU 5096 | AB-2017-11-29 | longitudinal | **D** (LSRII SORP, Aug 2017) |
| LAU 5022 | | (repeat) | |
| LAU 5099 | | BL only | |

= Panels used for tSNE analyses

Table S3. Staining reagents used for Flow Cytometry

| Target / marker | fluorochrome | Antibody clone | Source | Additional info (*) |
|---|---|---|---|---|
| A2/LLW | PE | NA | TCmetrix | HLA-A*0201 multimer with LLWNGPMAV (NS4b$^{214-222}$ from YFV) |
| | APC | NA | TCmetrix | |
| anti-rabbit IgG | PE | polyclonal (donkey) | eBioscience | secondary for TCF1: intracellular / intranuclear |
| | FITC | polyclonal (donkey) | eBioscience | secondary for TCF1: intracellular / intranuclear |
| CCR7 | Briliant Violet 421 | G043H7 | Biolegend | |
| CD127 | PerCP-Cy5.5 | A019D5 | Biolegend | |
| | Brilliant Violet 785 | A019D5 | Biolegend | |
| CD16 | Krome Orange | 3G8 | Beckman Coulter | |
| CD28 | A700 | CD28.2 | Biolegend | |
| CD38 | A700 | HIT2 | eBioscience | |
| CD45RA | ECD | 2H4LDH11LDB9 | Beckman Coulter | |
| | A700 | HI100 | Beckton Dickinson | |
| | Brilliant Violet 510 | HI100 | Biolegend | |
| CD58 | FITC | 1C3 | Beckton Dickinson | |
| CD58 | APC | TS2/9 | eBioscience | |
| CD69 | Brilliant Violet 650 | FN50 | Biolegend | |
| CD8 | APC-Alexa750 | B9.11 | Beckman Coulter | |
| CD8 | Krome Orange | B9.11 | Beckman Coulter | |
| CD95 | PC7 | DX2 | Biolegend | |
| CXCR5 | Brilliant Violet 605 | J252D4 | Biolegend | |
| HLA-DR | ECD | Immu-357 | Beckman Coulter | |
| | APC | L243 | Beckton Dickinson | |
| TCF1 | unconjugated | C63D9 (rabbit) | Cell signaling Technology | intracellular / intranuclear (primary) |
| PD1 | PerCP-eFluor710 | eBioJ105 | eBioscience | |
| | A488 | MIH4 | AbD serotec | |
| | BV711 | EH12-2H7 | Biolegend | |
| Fixable viability dye | Vivid Aqua (AmCyan) | NA | Invitrogen | |
| | Vivid Near InfraRed | NA | Invitrogen | |
| | eFluor 455 - UV | NA | Invitrogen | |

* unless otherwise stated, the antibodies were used in surface staining
